# Deep Learning-Based Prediction and Suppression of Protein Aggregation- Prone Regions

**DOI:** 10.1101/2024.03.06.583680

**Authors:** Vojtech Cima, Antonin Kunka, Joan Planas-Iglesias, Ekaterina Grakova, Martin Havlasek, Madhumalar Subramanian, Michal Beloch, Martin Marek, Katerina Slaninova, Jiri Damborsky, Zbynek Prokop, David Bednar, Jan Martinovic

**Affiliations:** IT4Innovations, VSB – Technical University of Ostrava, 17. Listopadu 2172/15, 70800, Ostrava-Poruba, Czech Republic; Loschmidt Laboratories, Department of Experimental Biology and RECETOX, Faculty of Science, Masaryk University, Kotlarska 2, Brno, Czech Republic; International Clinical Research Centre, St. Anne’s University Hospital, Pekarska 53, Brno, Czech Republic

**Keywords:** Amyloid, Aggregation, Deep Neural Network, Computational design, Solubility, Yield

## Abstract

Identification of aggregation-prone regions in proteins and their suppression through mutations is a powerful strategy to enhance protein solubility and yield, significantly expanding their application potential. Here, we developed a deep neural network-based predictor AggreProt, that generates a residue-level aggregation profile for protein sequences. The model outperformed or matched current state-of-the-art algorithms, as validated on two independent datasets comprising hexapeptides and full-length proteins with annotated aggregation-prone regions. We further validated the model experimentally using a set of 34 hexapeptides identified in the model protein haloalkane dehalogenase LinB, along with seven proteins from the AmyPro database. Experimental results agreed with our predictions in 79% of cases and also revealed inaccuracies in some database annotations. Finally, the algorithm’s utility was demonstrated by identifying aggregation-prone regions in the LinB enzyme and designing mutations to suppress aggregation in its exposed regions. The resulting variants exhibited reduced aggregation propensity, improved solubility, and up to a 100% increase in yield compared to the wild type. AggreProt is freely available to the scientific community via a user-friendly web server: https://loschmidt.chemi.muni.cz/aggreprot.

## 1. Introduction

Recombinant proteins are incredibly versatile with applications spanning from industrial biocatalysts, waste processing, the food industry, to highly efficient therapeutics or vaccines. The extraordinary advancements in computational structure prediction^1^ greatly accelerated their transformation towards one of the most promising technologies for a sustainable future and improved quality of life. Although protein structures can now be predicted with high accuracy almost instantaneously^2–4^, the underlying computational algorithms are designed assuming a single folded state corresponding to the global minimum of the free energy landscape^5^. In reality, folding of the protein chain into its functional - native state, driven by formation of intramolecular interactions and burial of hydrophobic residues within the protein core, competes with formation of non-native, intermolecular interactions leading to higher-order misfolded assemblies, i.e. protein aggregates. Misfolding pathways are inherent to any protein sequence and depend on both extrinsic (e.g., temperature, pH, ionic strength), and intrinsic (e.g., amino acid sequence, concentration) conditions^6^. Misfolding and aggregation can be triggered *in vivo* by numerous factors, such as oxidative stress, proteasome malfunctioning, posttranslational modifications, or mutations, and is a hallmark of severe pathologies such as Alzheimer’s disease, Parkinson’s disease, diabetes, prion disease, and many others^7^. Although the underlying mechanisms of these diseases are not yet fully clear, the cellular toxicity of the aggregates by both gain-of-function (generation of toxic species), and loss-of-function (i.e., reduction of proteins native activity due to their aggregation) have been described^8^. Moreover, aggregation and deposition of misfolded, insoluble proteins into inclusion bodies is one of the major hindrances in production of recombinant proteins in the biotechnology industry. Identification of sequence stretches driving the aggregation, i.e., aggregation-prone regions (APRs), is therefore of utmost importance.

Aggregation is inversely correlated with solubility, defined as the concentration of soluble protein in equilibrium with the solid aggregates. These can be classified according to their morphology to amorphous assemblies and amyloid fibrils or crystals. Amorphous aggregates lack well-defined high-order structure and their formation is driven by non-specific, mostly hydrophobic interactions^9,10^. Although analytical frameworks that quantitatively describe the kinetics of amorphous aggregation have been derived and successfully applied^11–20^, detailed characterization of amorphous aggregate assembly attracts little attention. Instead, amorphous aggregation is perceived as an omnipresent problem during protein production and processing^10^. In contrast, amyloids gained significant attention due to their link to many diseases^7^, and analysis of their formation has been extensively studied^21,22^. Amyloids are ordered two-dimensional assemblies formed by repeating units of soluble protein precursors (monomers) that are stabilised by an intermolecular network of hydrogen bonds within the cross-β-sheet architecture^23^. The interface within the amyloid subunit is formed by interdigitated complementary amino acid side-chains stabilised by van der Waals interactions, hydrophobic forces, and hydrogen bonds^24,25^. This structural and energetic complementarity in the eight possible arrangements (classes) of these so-called “steric zippers”^26^ governs the available amyloid sequence space^27^. Emergence of different fibril polymorphs, i.e., fibril structures with distinct morphologies formed by the same protein sequence, in the increasing number of experimentally determined fibril structures proves that the amyloid landscape is highly frustrated^28,29^. Despite the differences between chain arrangements within their cores, the polymorphs often share a common structural kernel which is believed to be an important driver of amyloid formation and essential for its stability^30–32^. These regions (APRs) are therefore perfect targets for designing mutations that decrease aggregation propensity and, consequently, increase protein solubility.

Existing algorithms for identification of APRs can be broadly categorised into (i) linear and (ii) structural predictors based on their underlying principle and input data. The first class derive the aggregation propensity from the protein sequence using the physio-chemical properties of the amino acids (e.g., hydrophobicity, net charge, patterning,), experimentally derived aggregation scales^33,34^, β- sheet propensity^35–39^, or their combination^40–44^. The second class of algorithms utilise protein structures to identify aggregation prone patches in globular proteins that are formed by sequentially distant residues that are impossible to identify using the linear predictors^45–48^. Emergence of databases compiling protein and peptide sequences with experimentally validated APRs or aggregation propensities^49–51^ boosted the development of machine learning-based predictors^27,52–59^. Their main advantage is that they are fast which (in some cases) enables their application on a proteome-wide scale^52^. However, most of them are only validated on existing datasets and their accuracy and usefulness in real-life applications are yet to be determined. The long-term evolution of aggregation predictors with distinct underlying principles greatly improved our understanding of protein aggregation on molecular level, and aided in solubilization and design of many important targets^60^. However, their performance remains suboptimal and there is still a great potential in improving their accuracy so that they can be efficiently incorporated into protein design pipelines to minimise the experimental screening effort.

Towards this, we have developed a novel protein aggregation predictor based on deep neural networks (DNN) called AggreProt. AggreProt consists of (i) sequential, and (ii) static models that were both trained on the hexapeptides from WaltzDB 2.0^51^ using atomic-related description of each amino acid (36 features^52^) of the hexapeptides or 18 features found in the database, respectively. The accuracy of AggreProt was higher or comparable to the state-of-the-art predictors Tango, Waltz, Aggrescan, PASTA 2.0, GAP, AnuPP, and FishAmyloid using both hexapeptide- and protein-based validation datasets. Furthermore, we carried out extensive experimental validation of AggreProt to demonstrate its application potential. In contrast to other predictors, AggreProt correctly identified APRs in one of our workhouse proteins - haloalkane dehalogenase (HLD) LinB (UniProt ID:D4Z2G1). This enabled us to fulfil our long-term effort and one of the motivations behind this study, i.e., to design mutations that improve the solubility and double the enzyme yield. We went beyond and experimentally validated AggreProt using hexapeptides derived from the LinB and proteins in the AmyPro database^50^. Based on our close investigation of the AmyPro entries using AggreProt, we have selected hexapeptides that were potentially misannotated in the database, and subsequently confirmed this by the experiment. This not only demonstrates the great accuracy and usefulness of the AggreProt, but implies that the benchmarking of aggregation predictors using AmyPro, CPAD 2.0 or similar datasets may result in biased statistical evaluation of the tools.

## 2. Materials and methods

### 2.1. Design and development of the aggregation prediction models

#### 2.1.1. Training and validation datasets

Training of the AggreProt DNN models was carried out using hexapeptides from the WaltzDB2.0 database^51^. WaltzDB contains 1416 sequences of hexapeptides with experimentally characterised and annotated amyloid-forming properties (515 amyloid, 901 non-amyloid). Additionally, each peptide entry contains features from three different in-silico predictors (TANGO^42^, PASTA 2.0^36^, and Waltz^40^) together with precalculated hydrophobicity and secondary structure propensities, and local energetic assessment from FoldX^61^. The hexapeptide dataset was randomly split into training (90%, WaltzDB-90) and validation (10%, WaltzDB-10) subsets. The training subset was used in a train-validation fashion (80:20 leave-one-out) during the hyperparameter optimization of the models, whilst the later subset of peptides was used for model evaluation.

The second validation dataset was selected from proteins in the AmyPro database^50^. AmyPro contains 162 protein sequences (AmyPro162) with manually annotated amyloidogenic regions collected from literature. To ensure there is no overlap between training and validation, proteins that contain sequences of the hexapeptides from WaltzDB2.0 were filtered out. In total, 37 proteins (Amy37) with annotated APRs remained and were used for validation. We further split the Amy37 into two subsets of proteins with APRs longer or shorter than 50 amino acids, and used the latter (n = 27 proteins, AmyPro27) as a high-confidence validation dataset. The number of sequences with short APRs in the complete dataset is 140 (AmyPro140). The main rationale behind such a split is that the annotations are often made based on indirect experimental evidence (as discussed in the results section) and true APRs are generally short segments.

#### 2.1.2. Metrics

The evaluation of DNN performance employed standard metrics derived from the confusion matrix, using both WaltzDB-90 in the training procedure and WaltzDB-10 in the testing (see below). True Positives (TP) are instances where ground truth positives (amyloid prone) are correctly predicted as positives, while False Positives (FP) refer to ground truth negatives (non-amyloid prone) erroneously predicted as positives. Conversely, False Negatives (FN) represent instances where ground truth positives are predicted as negatives, and True Negatives (TN) are instances where ground truth negatives are correctly predicted as negatives. Based on these definitions, Precision (Prec) was calculated as TP/(TP+FP), Recall (Rec) was calculated as TP/(TP+FN), and Specificity (Spec) was calculated as TN / (FP + TN). The performance assessment of the networks involved determining the Area under the Receiver Operating Characteristic (ROC) and Precision/Recall (PR) Curves (AuROCC and AuPRC, respectively). Youden’s J statistic^62^ was exploited on ROC curves to identify the optimal threshold for prediction.

The evaluation of the final models on whole protein sequence data (AmyPro) was done using the aforementioned metric and the Segment OVerlap score (SOV). SOV was defined by Rost and co-workers^63^, and we used the metric to evaluate the correctness of the predictions in a whole sequence context as it was used in ANuPP^52^. The SOV score can be calculated both for APR and non-APR segments. Final SOV values are given as the arithmetic average of individual SOV values calculated for each protein in the evaluated set (AmyPro37, AmyPro27) using the AggreProt optimal prediction threshold defined by Youden’s J.d.

#### 2.1.3. Exploring network attributes and hyperparameters. Architecture of the predicted models

AggreProt DNN models assess the aggregation tendency of an input hexapeptide by analysing either the 36 atomic features associated with each amino acid in the hexapeptide (linear model), following the approach employed in ANuPP^52^, or the 18 structural related features obtained from WaltzDB 2.0 database^51^ (static model). Particularly, the features considered by the static model are: scores from Waltz, TANGO, PASTA2.0 (parallel and antiparallel beta-sheet propensity), hydrophobicity (filter on the percentage of hydrophobic residues, including A, C, F, I, L, M, P, V, W, and Y), Chou-Fasman helix and strand propensities, and 11 FoldX force-field energy parameters calculated using Cordax^27^ Consequently, the initial input layer dimension (number of neurons) aligns with the count of initial features, which for the linear predictor is 36 per reside in the input hexapeptide, and for the static one is 18.

Throughout the training process, we fine-tuned the architecture and internal parameters of the networks using a 5-fold cross-validation with WaltzDB-90. We optimised the following parameters: i) number of bisectional layers (one or two), ii) number of dense layers (one, two or three), iii) dropout values (from 0 to 0.8, in 0.2-span steps), iv) number of neurons per layer (from 8 to 512, in powers of two-span steps), v) batch size (16 or 32), and vi) learning rate (from 0.0001 to 0.1, in powers of ten-span steps). The models were trained exploiting the TensorFlow Framework^64^ in our custom codebase.

A total of 840 and 1680 hyperparameter combinations were examined for the linear and static model architectures, respectively. Each of such combinations generated a unique network architecture that was trained in a 5-fold cross-validation manner. Thus, we generated 5 network instances for each architecture assessed. The performance of each architecture was assessed in terms of AuROCC. To avoid overfitting, the training was stopped at its optimum, when the training and validation ROC curves diverged (**Supplementary Figure 1**). As a final model we selected an average ensemble of the different network instances of the top performing architecture for each of the model types (linear and static).

Our models were trained on hexapeptide data. In order to make them amenable to process whole protein sequences, the input protein sequence of *n* amino-acids is divided into *n* hexapeptides using a sliding window approach with a step of one residue. The aggregation propensity prediction is generated for each hexapeptide. And all the information on a single amino-acid (contributed by up to 6 different hexapeptides) was aggregated using three different metrics: minimum, average and maximum.

#### 2.1.4. Model validation and benchmarking

The final models were evaluated using two different testing sets. First, the remaining 10% of WatzDB not used during the training procedure (WaltzDB10). Here, we calculated AuROCC as we did during training and compared to the training value.

Second, we validated on complete sequence proteins from AmyPro. To identify APRs on input protein sequences, these were fragmented into overlapping hexapeptides using a six-amino-acid- windows shifted one amino acid at a time. Each hexapeptide was evaluated by each of our two models (linear and static) and the results were averaged per residue mirroring the sliding window procedure used before. The performance of each network was assessed in terms of AuROCC, AuPR, and SOV.

### 2.2. Predictor testing and computational design of mutations suppressing the aggregation

#### 2.2.1. Detection of aggregation prone regions in model protein and computational design of aggregation-suppressing mutations

*Sphingobium japonicum* haloalkane dehalogenase LinB (UniProt ID: D4Z2G1) was analysed using the sequential model of AggreProt. The surface accessible area (SASA) was calculated using the DSSP algorithm^65^ on the publicly available LinB structure (PDB IDs: 1mj5). Residue accessibility values were normalized by the amino acid total volume to obtain SASA value per residue. Altogether, seven sequence stretches with highest AggreProt values were identified as potential APRs. The most solvent exposed residue with the highest AggreProt score in the middle of each APR, and two exposed residues that defined the APR borders were considered for mutagenesis. We iteratively mutated each selected position (APRs centres) or pairs of positions (APRs borders) to all possible amino acids and calculated the resulting AggreProt scores. Next, we evaluated the effects of all possible mutations on the structural integrity of the protein using HotSpot Wizard (HS)^66^ and Rossetta ddg_monomer^67^. The mutations that had low AggreProt score, high mutability according to HS, reasonable occurrence (i.e., not rare) in homologous sequences based on multiple sequence alignment, and did not significantly compromise the conformational stability of the protein, were selected for experimental characterization.

#### 2.2.2. Identification of hexapeptides for experimental validation of their aggregation propensity

Based on the manual inspection of our AmyPro37 validation dataset, we selected seven relatively short proteins (lysozyme c, apomyoglobin, serum amyloid A, envelope small membrane protein, cell division topological specificity factor, gallectin-7, and gamma-crystallin D) with clear deviations between our predictions and ground truth based on AmyPro. Within these proteins, we identified 17 hexapeptides which might be incorrectly labelled in the AmyPro as amyloids (i.e., false positives - FP, n = 11) or non-amyloids (i.e., false negatives - FN, n = 6). Our selection criteria included inconsistency between labels and primary literature, and low or high AggreProt scores for FP and FN, respectively. As a second dataset, we chose 7 hexapeptides corresponding to the putative APRs found in LinB and their 10 counterparts containing aggregation-suppressing mutations described above. Altogether, we selected 34 hexapeptides for experimental validation of their aggregation propensity (**Supplementary Table 1**).

### 2.3. Experimental validation

#### 2.3.1. Materials and chemicals

The genes encoding LinB variants with C-terminal His-tag were cloned in the pET21b plasmids and expressed in *E. coli* BL21(DE3) (New England Biolabs, UK). Target mutations were introduced using megaprimer PCR mutagenesis using pairs of mutagenic and non-mutagenic primers (**Table 1**). All chemicals used in this study were purchased from Sigma (USA) unless specified otherwise. The hexapeptides selected for the experimental validation were synthesised by ABI Scientific Inc. (USA) as the trifluoroacetate salts with acetylated and aminated N- and C-termini, respectively. Their purity was 98 % according to the HPLC analysis carried out by the manufacturer.

**Table 1:**
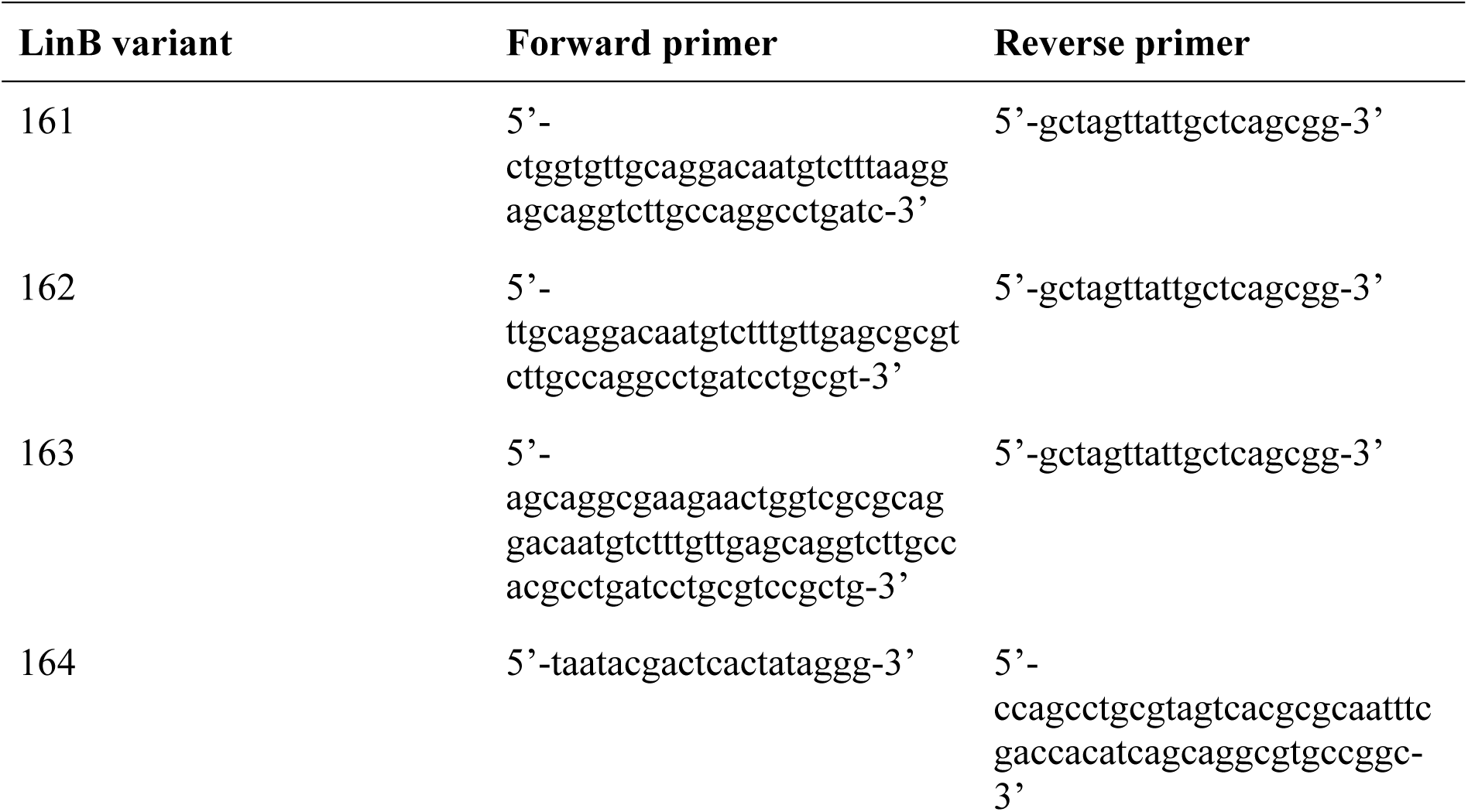
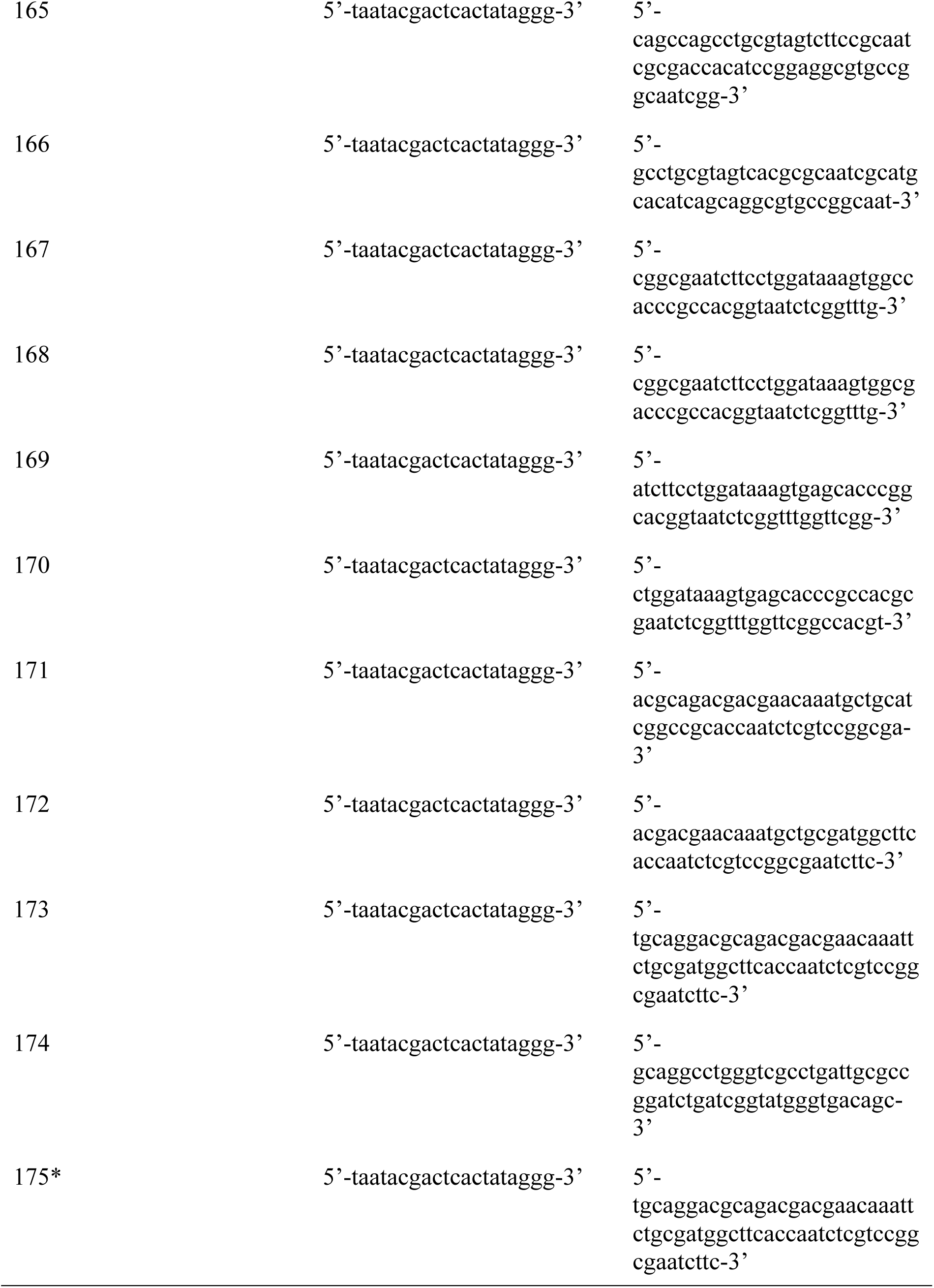
List of mutagenic primers. LinB175 (denoted with an asterisk) was prepared using the DNA sequence of LinB165 as a template. All other mutants were prepared using wild-type as a template.

**Table 1:**
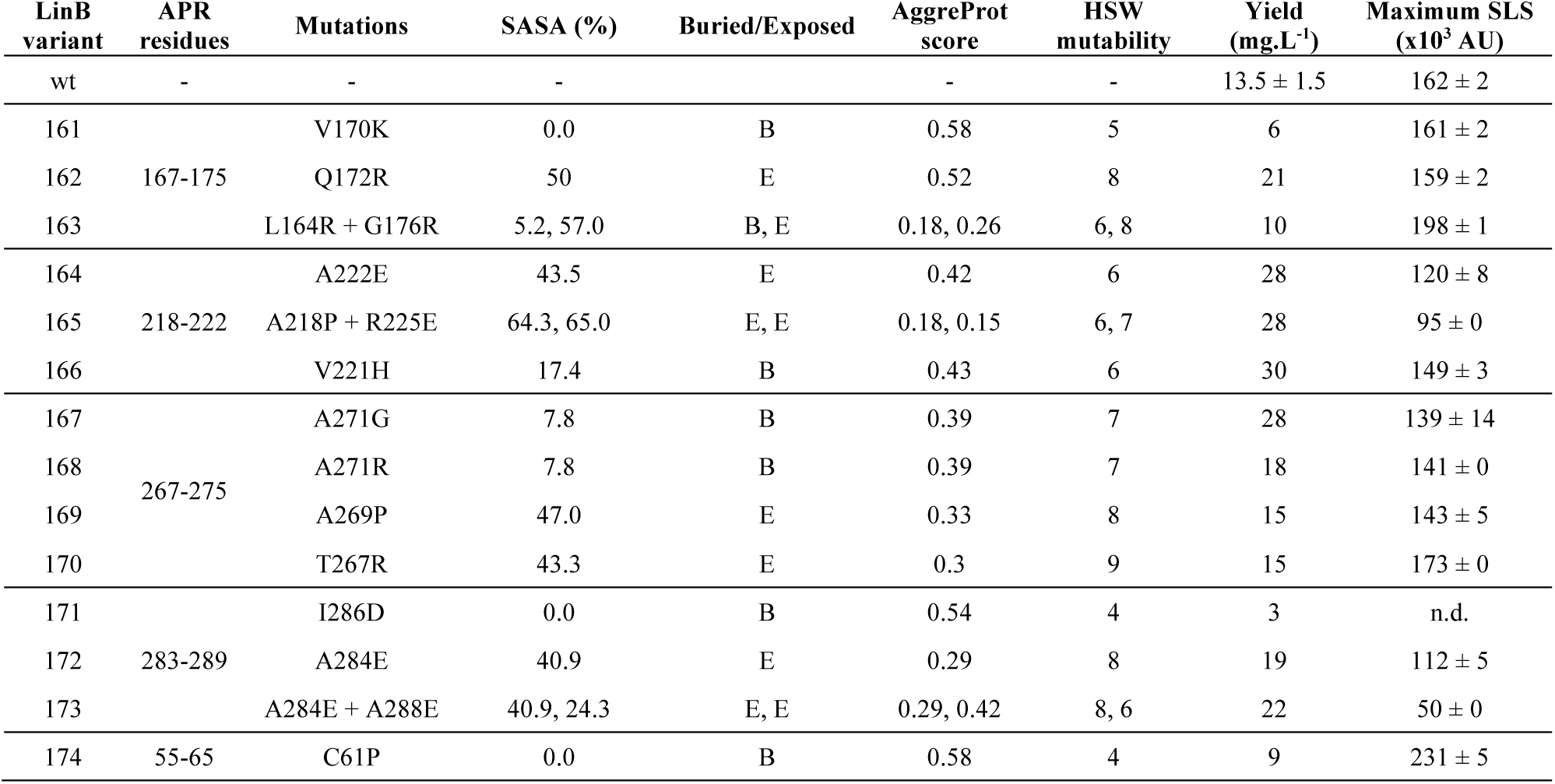
Design of mutations to the APRs that suppress aggregation. The SASA (solvent-accessible surface area) has been calculated from the structure (PDB-ID:1MJ5) using DSSP^65,76^ and ranges from 0 (completely buried - B) to 100 (completely exposed - E). HotSpot Wizards (HSW) score provides mutability of a given residue based on the multiple sequence alignment of homologous sequences and ranges from 0 (not mutable) to 10 (mutable). Yield of the soluble monomeric protein from 1 L of bacterial culture. The value given for wild type is the mean and standard deviation between two individual batches. Maximum SLS corresponds to the highest values experimentally measured during melting scan from 15 to 95 °C at 1 °C/min scan rate at 1 mg.ml^-1^ of protein.

#### 2.3.2. Site-directed mutagenesis

The mutagenic primers used for the introduction of the designed mutation were ordered from Merck (Merck, Germany). The results of the megaprimer mutagenesis, consisting of two-round PCR, were verified using sequencing (Eurofins Genomics, Germany).

#### 2.3.3. Protein expression, purification, and quality control

LinB variants were expressed in *E. coli* BL21 (DE3) transformed by the mutagenic plasmids (pET21b) encoding respective LinB mutants. The protein expression was induced by the addition of IPTG (0.4 mM final concentration) and carried out at 20°C for 16 hours. The cells were harvested by centrifugation (4,000 x g, 4 °C, 30 min) and frozen at -70 °C. After defrosting, the resuspended cells were disrupted by the addition of DNAse I and 3 rounds of sonication. The disrupted cells were centrifuged (21 000 *g*, 4 °C, 60 min) and the proteins were purified from the cell lysates by metallo- affinity chromatography with HisTrap HP column (Cytiva, USA) charged with N^i2+^ ions using the stepwise increase of imidazole in elution buffer. The pure monomeric fraction for the following aggregation experiments was isolated by size-exclusion chromatography using the Superdex 200 pg 10/300 GL column (Cytiva, USA) with the 50mM Potassium phosphate buffer (pH 7.5) used as the mobile phase. Protein purity was verified by SDS-PAGE, and the concentration determined by UV-absorbance measurement the DS-11 Spectrophotometer (DeNo-vix,USA).

#### 2.3.4. Temperature scanning experiments

The aggregation propensity of all tested LinB variants was screened using UNcle (Unchained labs, USA) in a 50 mM potassium phosphate buffer pH 7.5. The protein samples were continuously heated from 20 to 95 °C at 1°C/min scan rate and the static light scattering (SLS) at 266 nm and fluorescence emission spectra were recorded. The aggregation propensity was estimated from the maximum amplitude of the SLS curve, while the apparent melting temperature (*T*m^app^) was evaluated from the midpoint of the average emission wavelength (i.e., Barycentric mean - BCM) curve. Each sample was analysed in a duplicate at a protein concentration of 0.275 mg/ml to ensure comparability between individual variants.

#### 2.3.5. Aggregation kinetics

Several LinB variants were selected based on the previous screening and their aggregation kinetics at 37 °C were monitored using SLS and Thioflavin T assay (ThT) in 50 mM Potassium phosphate buffer (pH 7.5). The scattering intensity at 266 nm was measured using UNcle (Unchained labs, USA) and the amount of aggregates was estimated from the curve amplitude. Each sample was analysed in a duplicate at two different protein concentrations of 0.5 and 1.0 mg/ml.

The ThT assay was measured in 384-well plates (Corning® 384-well black, clear flat bottom polystyrene microplate) sealed with aluminium tape (Corning® 96 Well Microplate Aluminum Sealing Tape in duplicates using the Synergy H4 reader (BioTek, USA). The ThT fluorescence (30 µM) added to samples was excited at 440 nm and emission measured at 485 nm using bottom read over the course of 24 hours in 5-minute intervals at 37°C. Each variant was measured at two protein concentrations of 0.5 and 1.0 mg/ml in duplicates.

#### 2.3.6. Flow-induced dispersion analysis

The amount of soluble protein in the samples after aggregation was analysed for the two least aggregation-prone LinB variants and wild-type using flow-induced dispersion analysis (FIDA) using FIDA 1 (Fidabio, Denmark) equipped with the protein intrinsic fluorescence detection (280 nm excitation). The three proteins (1 mg/ml) were incubated at 37 °C for 24 hours in low binding tubes. Aliquots withdrawn from the aggregation reaction at the start of incubation (0 h), and after 1 and 24 hours were centrifuged (18000g, 4 °C, 20 min) and the supernatant analysed using FIDA with the following method: Step 1: NaOH wash (60 s, 3500 mbar), Step 2: H_2_O wash (60 sec, 3500 mbar), Step 3: Buffer wash (40 sec, 1500 mbar), Step 4: Sample application (20 s, 75 mbar), Step 5: Sample analysis (buffer, 75 s, 1500 mbar). The amount of soluble protein at each time-point was determined from the integral of the resulting Gaussian peak and expressed relative to the amount at the beginning of the reaction.

#### 2.3.7. Dehalogenase activity measurement

Effects of the introduced mutations to the protein functionality were verified by determination of their specific activity towards 1,2-dibromoethane using the colorimetric assay according to Iwasaki^68^. The dehalogenation reactions were carried out at 37 °C in 25-ml ReactiFlasks closed by Mininert Valves. The reaction mixture was composed of 10 mL of 100 mM glycine buffer (pH 8.6) and 10 μl of the substrate (1,2-dibromoethane, purity ≥ 98%). The reaction was initiated by the addition of the enzyme (final concentration ∼ 2 µg/ml). The progress of the reaction was monitored over 30 minutes by withdrawing 1 ml aliquots from the reaction mixture in 4-minute intervals. At each time point, the reaction was terminated by the addition of 0.1 ml of 35% nitric acid to the mixture. The bromides formed during the reaction were detected spectrophotometrically upon the reaction with mercuric thiocyanate and ferric ammonium sulphate using absorbance at 460 nm using the Eon plate spectrophotometer (BioTek, USA). The absorbance read-out was recalculated to the concentration of the product using the calibration curve with known concentration of bromides. The activity was determined from the slope of the product concentration over time.

#### 2.3.8. Circular dichroism spectroscopy

The secondary structure of the two least aggregation-prone LinB variants and wild-type and their thermal stability were analysed using circular dichroism (CD) spectroscopy. The CD spectra of 0.175 mg/ml proteins in a 50 mM potassium phosphate buffer were measured from 185 to 260 nm (0.25 s integration time, 1 nm bandwidth) at 20 °C using the Chirascan spectropolarimeter (Applied Photophysics, UK). Thermal stability was probed by monitoring the ellipticity changes at 227 nm while the temperature was gradually increased at 1°C/min scan rate from 20 to 75 °C. The apparent melting temperatures were assessed from the midpoints of the denaturation curves.

#### 2.3.9. Thioflavin T assay and transmission electron microscopy of the hexapeptide aggregates

The peptide lyophilisates were first dissolved in a 50 mM sodium phosphate buffer (pH 7.4) to a final concentration of 5 mM. The insoluble ones (based on turbidity and physical appearance of the buffer stocks) were additionally dissolved in either DMSO or HFIP (for peptides containing cysteine to avoid their oxidation) to a final concentration of ∼100 mM. For the aggregation assays, all peptides were diluted to the final working concentration of 1 mM using 50 mM sodium phosphate buffer pH 7.4 containing 0.05% sodium azide to prevent bacterial contamination. Samples were incubated at 37 °C for 4 weeks and the progress of aggregation was monitored using ThT assay as described above. At the end of the incubation, full ThT emission spectrum was recorded (ex. 440, em. 460-560 nm) for each sample to determine whether the hexapeptide aggregate is positive for ThT. Finally, 4 µl of the 1 mM peptide solution was spotted onto a carbon-coated 300 mesh copper grid and incubated for 2 minutes, followed by 2 minute negative staining using 2% uranyl acetate solution. Imaging was performed using Talos™ F200C microscope (Thermo Scientific™). For reliable comparison, three independent regions of interest were imaged for all peptides and the obtained structures were characterised into five different categories: fibrils, short fibrils, amorphous aggregates, hydrogels/unspecific structures and grids with no aggregation. Classification of the hexapeptide states to amyloids and non-amyloids for AggreProt benchmarking was done primarily based on the TEM images by the experimentalist without knowledge of the predicted aggregation scores of the hexapeptides to avoid biassed assessment.

## 3. Results and Discussion

### 3.1. Derivation of the input features for the training dataset and architecture of the deep neural networks

One of the major hindrances in the development of reliable protein aggregation predictors is the scarcity of reliable experimental data for training. We have identified AmyPro^50^ and WaltzDB2.0^51^ as the most suitable data repositories. AmyPro contains 162 protein sequences with manually annotated amyloid forming sequence stretches (APRs), whereas WaltzDB2.0 involves 1416 hexapeptides with experimentally verified amyloid-forming propensities. Considering the unambiguous data labels in the later, and the fact that the hexapeptide is widely considered as the minimal structural unit capable of forming the steric zipper of the amyloid core^23^, we have decided to utilise the set of hexapeptides from WaltzDB for the training.

In the next step, we have selected three classes of descriptors for the hexapeptides. For the first model class (termed “sequential”), input features for every amino-acid in each hexapeptide were generated using the amino-acid atomic composition (36 descriptors^52^). For the second class (termed “static”), we have used the parameters from the WaltzDB itself. These include aggregation propensity scores calculated using Waltz, TANGO, and PASTA software, Chou-Fasman alpha-helix and beta-sheet propensity^69^, hydrophobicity scale (as described in WaltzDB^51^), and eleven energy parameters calculated using FoldX^61^ and Cordax^27^.

The two models were trained on a 5-fold cross-validation procedure (see Methods), diverged on the type of input features (see above), and both took a hexapeptide as input. From the training procedure, we chose the top-performing architectures (summarized in **Figure 1**) and we built each model as an ensemble of the 5 variations of each top-ranking architecture produced by the 5-fold cross-validation procedure. The total number of parameters incorporated into the networks was 46113 and 2049 for the linear and static model, respectively.

**Figure 1.**
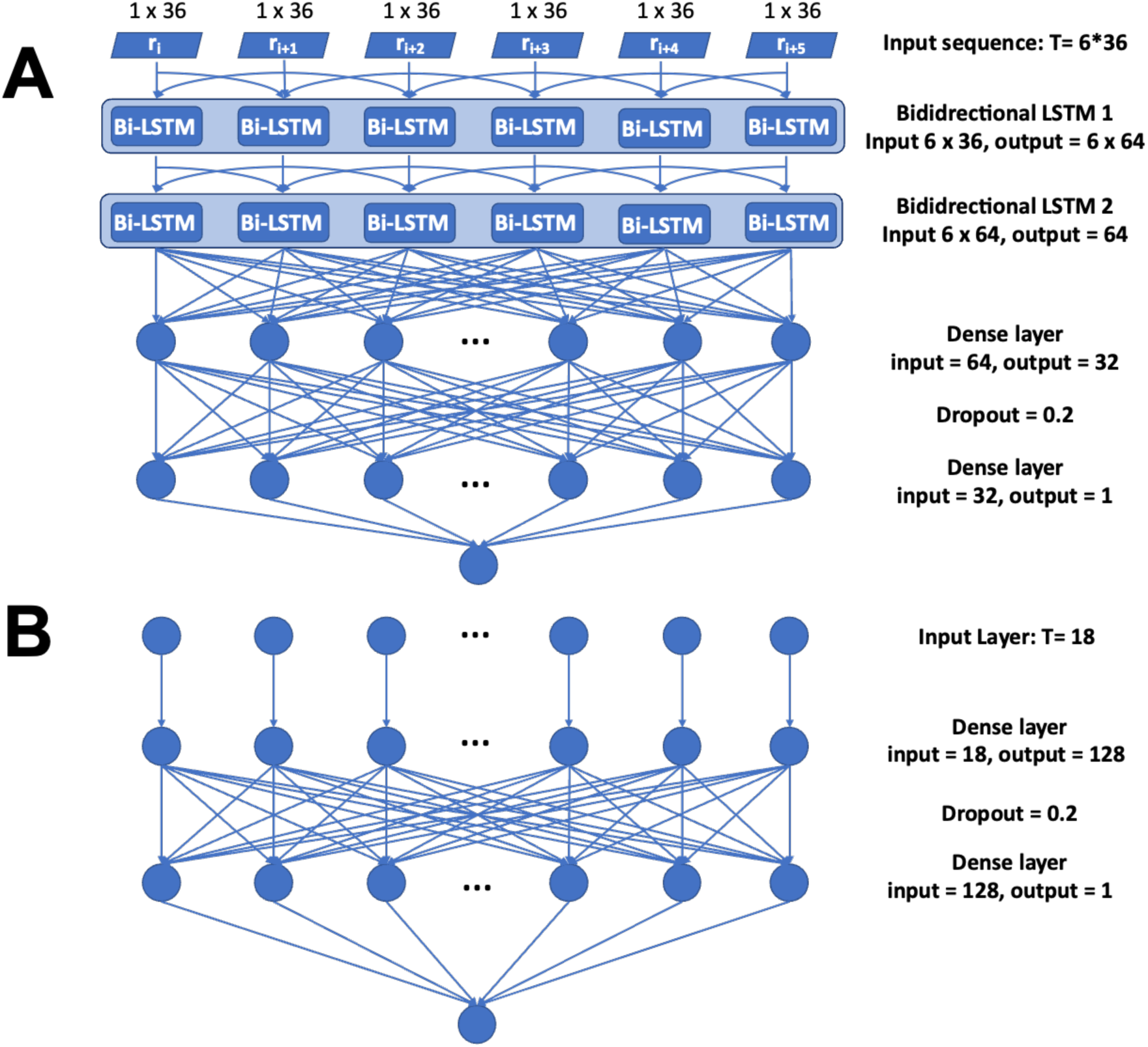
Model architectures: **A** In the linear model, an input layer processes sequential inputs into a cascade of bidirectional long short-term memory units, followed by two dense (fully-connected) layers, each connected to their corresponding dropout layer, finishing in a single-unit dense output layer. **B** The static model is composed of an input layer that processes normalized static features (hexapeptide-level properties) into a dense (fully connected) layer, followed by a dropout layer and a single-unit dense output layer. All model architectures are supported with by dropout layers aiming to reduce potential model overfitting (**Supplementary Figure 1**).

### 3.2. Model validation and benchmarking

We validated our models against two different datasets. Initially, we used a dataset containing the remaining 10% of WaltzDB hexapeptides (WlatzDB-10) not used during the training-validation procedure. The linear and static models achieved AuROCC of 0.89 and 0.85, respectively, which is comparable to the values obtained during the training (0.88 and 0.89). Both models reached AuROCC of 1 (**Supplementary Figure 1**) on the training set, altogether confirming that overfitting and data leakage were avoided.

Second, we enquired how our models performed on complete protein sequences. Here, we implemented a two-step sliding window procedure to first decompose the input protein into hexapeptides and then map and aggregate each hexapeptide prediction (value) to every amino acid in the protein sequence (see Methods). We used three different aggregation metrics (minimum, average, and maximum) to map the hexapeptide prediction onto individual amino acids. On a dataset completely oblivious to any hexapeptide used during the training validation procedure (AmyPro37), the linear predictor achieved an AuROCC of 0.53 for all three aggregating metrics, and an AuPRC of 0.23. The static predictor reached an AuROCC ranging between 0.49 (minimum) and 0.50 (average and maximum); all three metrics resulted in an AuPRC of 0.23 (**Supplementary Figure 2 A, B**).

The striking differences between the validation results on the whole-sequence and hexapeptide-level data could be rationalized by the closer inspection of the AmyPro database. We observed that AmyPro annotations often include unusually long APRs (more than 50 consecutive residues). It is highly unlikely that all residues in these stretches contribute equally to the aggregation propensity of the protein, which is not reflected by the binary annotation of the database (i.e., amyloid, non-amyloid) and can thus introduce bias during validation. Removing proteins containing such long APRs from the previous dataset, we obtained AmyPro27, and we proceeded again to validate our models. Using the curated dataset, the linear model reached AuROCCs of 0.59, 0.64 and 0.65 for each of the minimum, average, and maximum aggregation metrics, respectively. The corresponding AuPRCs were 0.28, 0.32 and 0.33. The static predictor achieved AuROCCs of 0.61, 0.66, and 0.65 for the three aggregation matrices, and their corresponding AuPRCs were 0.32, 0.35 and 0.34. (**Figure 2 A, B**) This confirms that our predictor may not be suitable for the detection of long APRs, whose biological relevance is discussed further in the text.

**Figure 2.**
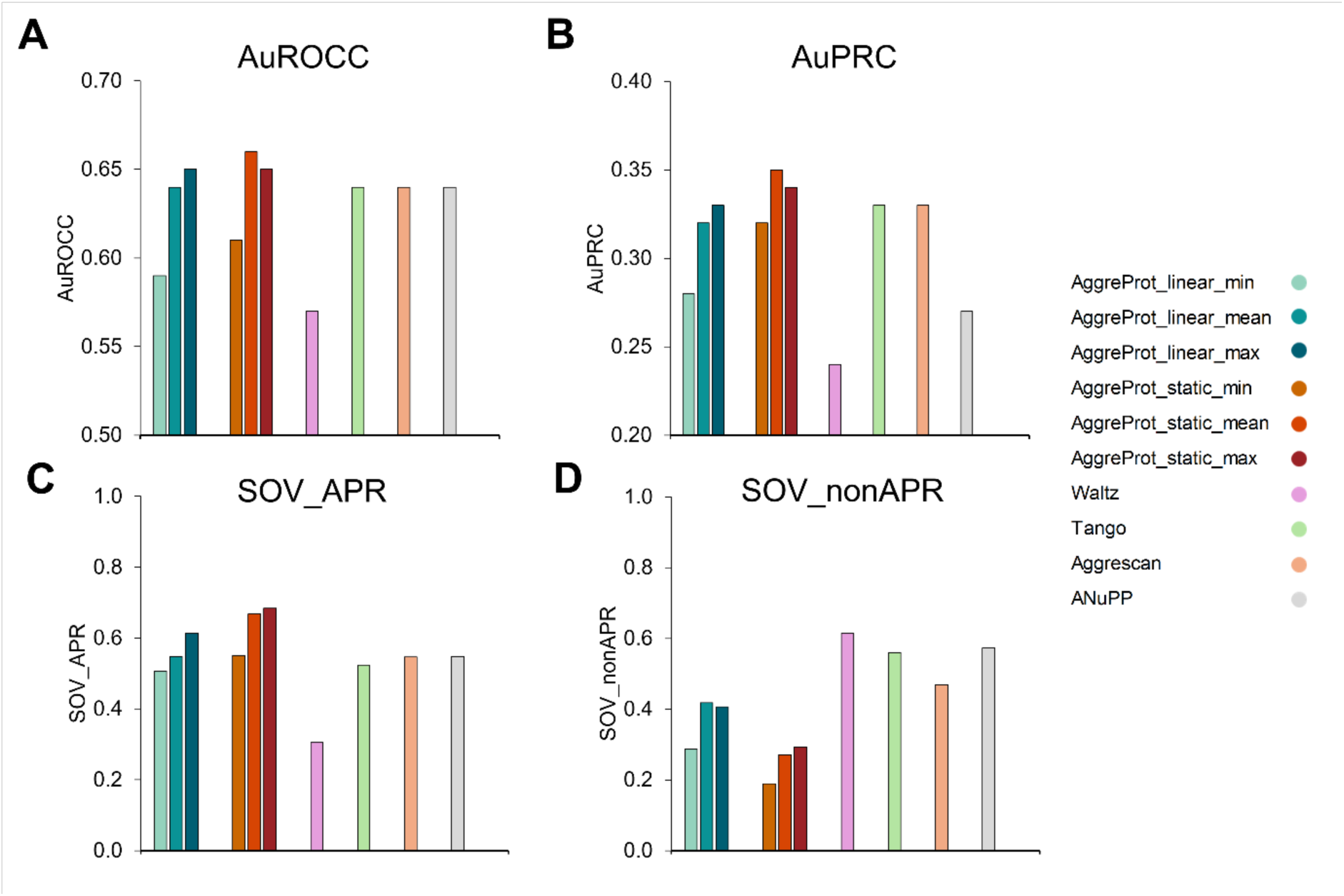
Performance of the trained networks using different aggregating metrics and their comparison to the state-of the art predictors on AmyPro27. Different evaluation metrics are shown for the performance of the linear (blue hues) and the static (red hues) predictors on AmyPro27, along with Waltz (pink), Tango(green), Aggrescan (orange), and ANuPP (grey). **A**: Area under the Receiver Operating Characteristic Curve (AuROCC). **B**: Area under the Precision-Recall Curve (AuPRC). **C**: Averaged Segment OVerlap score for Aggregation Prone Regions (SOV_APR). **D**: Averaged Segment OVerlap score for Non-Aggregation Prone Regions (SOV_nonAPR). The magnitude of each metric is shown in the y axis.

Third, we computed the SOV metric, which represents how well our predictions cover the APRs and non-APRs of the annotated proteins in AmyPro (in a range between 0 and 1). The results shown herein represent the arithmetic average over all proteins in the evaluated set. The different aggregation metrics on the linear predictor ranged between 0.51 and 0.61 in APRs, and between 0.29 and 0.42 in non-APRs when validated on AmyPro27, while in the static model, they ranged between 0.51 and 0.68 in APRs and between 0.19 and 0.29 in non-APRs (**Figure 2 C, D**). Using AmyPro37, these values dropped to a range between 0.40 and 0.47 for APRs and a range between 0.23 and 0.34 for non-APRs, in the case of the linear predictor, and to values between 0.48 and 0.53 for APRs and between 0.15 and 0.23 for non- APRs in the case of the static predictor (**Supplementary Figure 2 C, D**).

In general, we didn’t observe significant differences between the performance of the linear and static predictors. The latter performed better based on some aggregation metrics, but the improvement was outweighed by the larger computational time. Based on these results, we selected the linear model with an average aggregation procedure (AggreProt_linear_mean) and an optimal threshold of 0.25 calculated by Youden’s J^62^ on AmyPro140 (the larger validation dataset available without long APRs) for predictions described further in the manuscript. The same predictor has been implemented in AggreProt web server^70^, where details of its validation are also shown. Finally, we compared the results of our models to those of other sequence-based state-of-the-art methods, namely Waltz^40^, Tango^42^, ANuPP^52^, and AGGRESCAN^33^. Except in the case of SOV for non- APRs, our models performed equal to or better than the others (**Figure 2**, **Supplementary Figure 2**). The underperformance on the SOV metric for non-APRs indicates that our networks estimate aggregation-prone regions that are not annotated as such. Based on our experimental evaluation (described in detail in section 3.4) this might stem from a poor data annotation rather than being a model performance issue.

### 3.3. Introduction of mutations to the APRs identified by AggreProt reduces aggregation and increases the solubility of LinB

Next, we tested our newly trained predictor on a model haloalkane dehalogenase LinB. HLDs (EC3.8.1.5) are well-characterised model enzymes belonging to a larger superfamily of α/β hydrolases that hydrolytically cleave carbon-halogen bonds in a broad range of aliphatic halogenated compounds. LinB has been isolated from the bacterium *Sphingobium japonicum UT26* and its ability to convert persistent manmade pollutants and xenobiotics, most notably hexachlorocyclohexane, makes it a valuable target for industrial biodegradation or bioremediation^71–73^. Its broader and efficient application is, however, hindered by its poor solubility and low expression yield compared to the rest of structurally similar HLDs. The enzyme was previously stabilised by approx. 23 °C in terms of apparent melting temperature using a combination of energy calculations, disulfide bridge design, and molecular dynamics simulations^74^. However, the remarkable increase in its thermostability has not been accompanied by improvement in yield and solubility, suggesting distinct determinants of LinB’s conformational and colloidal stability. Our previous efforts to identify APRs in LinB by the state-of-the-art predictors and design solubilizing mutations have been futile.

Using the linear model of AggreProt we identified 7 regions within the LinB sequence whose score was higher compared to the rest of the sequence and could thus represent potential APRs (**Figure 3A**). We defined each APR empirically as any continuous residue stretch with a minimum length of 6 amino acids containing a maximum of 1 residue below the aggregation threshold (0.25, see above). Two of these APRs were also consensually detected using other aggregation prediction algorithms, whereas the others were identified only by a few or none (**Supplementary Figure 3**). We immediately excluded the three stretches towards the N-terminus since they are buried within the central beta-sheet of the hydrolase core (**Figure 3A**, grey APRs), and focused on the remaining four on the protein surface. Next, we selected residues that were near the centre or the edges of the APR and evaluated changes in aggregation score upon their *in silico* saturation mutagenesis. We purposely selected APR residues with different degrees of solvent exposure to evaluate the importance of SASA for the design of aggregation-suppressing mutations. The five residues that lowered the aggregation the most regardless of the APR according to AggreProt were R, K, P, E, and D, which are known to be the "aggregation gatekeepers"^75^. Conversely, hydrophobic residues such as V, I, or C were among the worst, as expected. We further analysed the proposed mutations by HotSpot Wizard^66^ and Rosetta^67^ to assess their mutability and effect on conformational stability, respectively. Finally, we selected 3-4 mutations from each APR, yielding 14 LinB variants for experimental validation.

**Figure 3:**
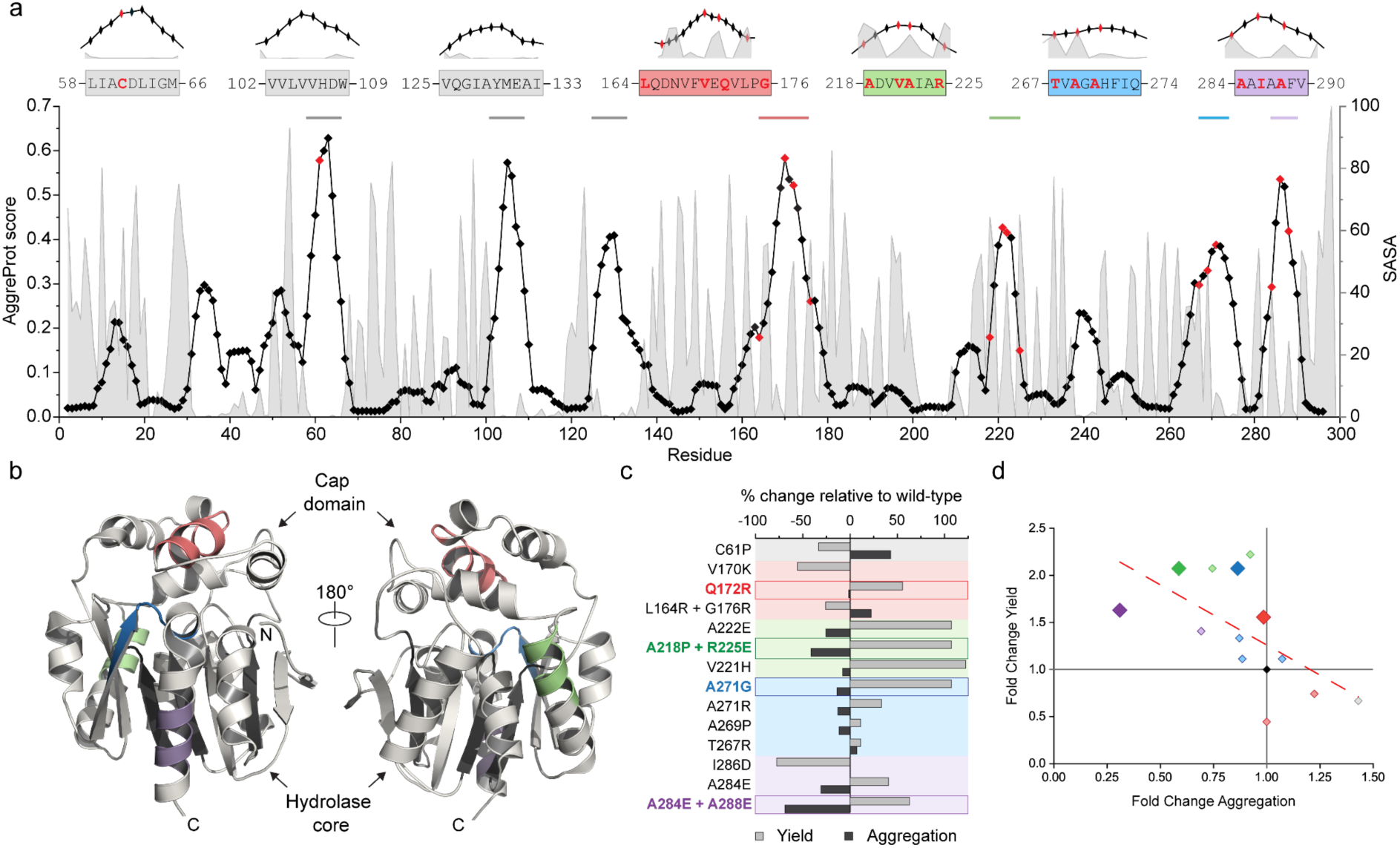
Identification of APRs in the model haloalkane dehalogenase LinB and their suppression by computational mutation design. **a) Aggregation profile of LinB sequence**. The AggreProt score for each LinB residue (black line and symbols) is overlaid with the relative SASA score (grey areas). Residues selected for mutagenesis are marked in red. The zoom-ins of the seven regions with the highest aggregation score are shown on top. The APRs1, 2, 3, and 4 are coloured red, green, blue, and violet, respectively. **b) Structural context of the APRs** identified by AggreProt in LinB (Pdb-ID: 1MJ5).. **c) Changes in aggregation propensity and yield induced by the designed mutations.** Total yield is calculated from the amount of soluble monomer isolated using SEC. Aggregation score is taken as the maximum value of the SLS signal during melting scans of the variants at 0.275 mg/ml and 1°C/min scan rate. The relative scores are calculated as wt - wt/mut. The mutations subjected to subsequent characterization are colour-coded according to the respective APR from a and highlighted with the box. **D) Correlation between solubility and aggregation.** Relative scores from c are plotted and their correlation is depicted by the red dashed line. The variants selected for further characterization are highlighted.

We expressed all proteins in two batches of seven mutants with wild type in each, and purified them from cell free extract using a combination of metallo-affinity and size-exclusion (SEC) chromatography. We managed to obtain all but one (I286D) variant to excellent purity (>95% based on SDS-PAGE) and homogeneity (>99% based on SEC). We evaluated the solubility of each variant from (i) the SDS-PAGE analysis of the total, soluble, and insoluble fractions during expression, and (ii) the final yield of monomeric protein. Unfortunately, the former strategy proved to be highly irreproducible with results of two large-scale and two small-scale expressions of wild-type alone differing by 80% (**Supplementary Figure 4**). In contrast, the deviation between yields from two large-scale expression and purification batches of wild type differed by only 11% so we used this parameter for comparing soluble expression of individual variants (**Table 1**). Notably, the expression and purification of all variants have been carried out over the course of two consecutive weeks by a single operator, using the same batches of media, incubation times, and purification protocol to ensure maximum comparability.

In total, eight out of fourteen variants had significantly improved yield, two mutations had a neutral effect, and four variants had lower yield compared to the wild type (**Figure 3c**, grey). The latter are mutations of the buried residues, confirming our hypothesis regarding the importance of SASA during design (**Table 1**). The biggest improvements in yield were achieved with mutations in the APR2 (green in **Figure 3**, residues 218-225), APR3 (blue, 267-274) and APR4 (purple, 284-290), confirming their correct identification by AggreProt. Arguably, APR1 has been identified correctly (judged by the Q172R, Table 1) but more efficient suppression of its effect will require further design effort. To assess the effect of the mutations on the aggregation propensity of LinB, we carried out a temperature melting scan and used static light scattering (SLS) as a measure of the aggregate formation (**Figure 3c**, black; **Supplementary Figure 5**). Similarly to yield, the mutations of buried residues increased aggregation (I286D, which introduces a buried negative charge, could not even be obtained in monomeric form). The rest of the mutations suppressed the aggregation or had a neutral effect (**Figure 3c**, black). We found a weak negative correlation (R^2^ = 0.36, Pearson’s Correlation Coefficient = -0.60) between changes in aggregation and soluble expression (**Figure 3d**), suggesting that these two properties are linked. On average, the introduction of negative charge (mutations to E or D) has a better effect on aggregation and yield compared to the positive charges (R or K). However, a systematic screening of more mutants is needed before broader generalisation can be drawn.

Finally, we picked the best mutant from each APR for subsequent validation (highlighted by a coloured box in **Figure 3c**). We monitored their aggregation kinetics at 37 °C for 24 hours using SLS and Thioflavin T (ThT), which is an established amyloid fluorescence probe (**Supplementary Figure 6A, B**). The aggregation kinetics are characterised by a fast increase of SLS signal followed by its slower attenuation and subsequent slow increase to a plateau. The ThT signal shows a short lag phase on a time scale corresponding to the fast phase of SLS signal, and is followed by an exponential increase to a final plateau. The multiphasic nature of both signals suggests a complex aggregation mechanism of LinB, whose description is beyond the scope of this work. However, the final SLS and ThT signals report well on the amount of aggregates in the samples (**Supplementary Figure 6C**). The SLS signals of the single point mutants in APR1 and 3 are ca. 50% higher compared to the wild type, suggesting that their aggregation propensities have not been significantly improved. In contrast, double-point mutants in both APR2 and APR4 show remarkably reduced aggregation based on both SLS and ThT. This is further supported by additional analysis of their soluble fraction upon centrifugation using flow-induced dispersion analysis (FIDA), which detects only 40% of residual monomer in the wild type sample, but 76 and 78% in case of LinB165 (A218P + R225E, APR2) and LinB173 (A284E + A288E, APR4), respectively (**Supplementary Figure 7**). This validates the results of our initial screening and confirms that the aggregation propensity of the designed LinB variants has been significantly decreased. Moreover, we further verified that the activity of both LinB variants towards 1,2-Dibromoethane has not been compromised by the mutations, which makes them perfect targets for future research and biotechnological applications (**Supplementary Figure 8**).

In summary, AggreProt identified four APRs in haloalkane dehalogenase LinB which were successfully suppressed by mutagenesis leading to LinB variants with increased solubility and yield, and decreased aggregation propensity.

### 3.4. Experimental validation on hexapeptides reveals inaccuracies in the annotations of aggregation databases

Finally, we experimentally validated the AggreProt predictions using a set of newly identified 34 hexapeptides. These included (i) 7 peptides corresponding to the APRs identified by AggreProt in LinBwt, (ii) 10 LinB-derived peptides carrying mutations aiming to suppress the aggregation described above, and (iii) 17 peptides selected from proteins in AmyPro database (**Supplementary Table 1**). The first two subsets were selected to test the performance of AggreProt on the identified APRs outside their structural context in LinB. Both TEM and ThT evidence were used to establish the classification of the hexapeptide states into amyloids and non-amyloids. However, TEM images were preferred to avoid human biased assessment of the difficult-to-interpret ThT experiments (**Supplementary Figure 9**). Amyloid fibrils were found in all 7 samples after 1-month incubation, confirming their accurate identification. Interestingly, aggregates were also observed in the samples of peptides carrying putative aggregation-suppressing mutations (**Figure 4b**). However, the amount of aggregates was generally lower and their morphology varied from amorphous clumps and hydrogels to short fibrils, indicating that the aggregation propensity of the peptides was reduced, but not completely suppressed by the mutations. The only peptides that showed absolutely no signs of aggregation were the two derived from LinB APR4 (corresponding to LinB167 and 168). The results obtained with these two subsets of hexapeptides, together with the model case above, illustrate that although AggreProt was successful in the identification of APRs, the design of the suppressing mutations requires further optimization.

**Figure 4:**
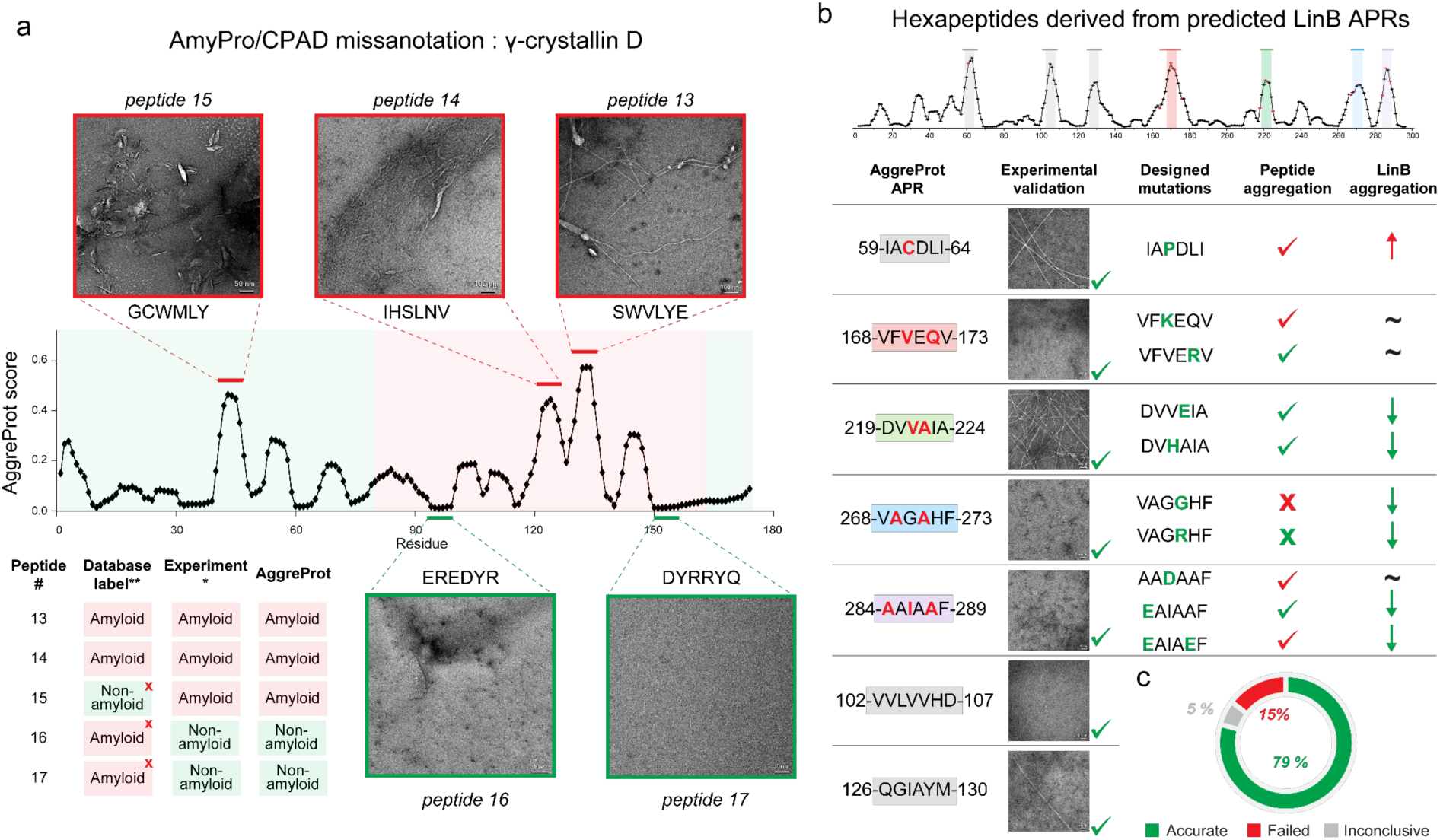
Experimental validation of AggreProt using hexapeptides identified in proteins from AmyPro database and LinB. a) Example of hexapeptide selection using gamma-crystallin D entry from AmyPro (CPAD 2.0) database. AggreProt score (black line and points) is not in agreement with the database annotation of amyloid (red shading) and non-amyloid (green shading) regions. Five hexapeptides were selected to represent potentially false negative (peptide 15; red), false positive (peptides 16, 17; green), and true positive (peptides 13, 14) amyloid annotation. Experimental validation by transmission electron microscopy (TEM, insets; red) and Thioflavin T assay confirmed that the AggreProt predictions are correct and three of the respective segments are misannotated in the databases (sequences corresponding to peptides 15-17). Similar cases of misannotations were found in other proteins in the databases (**Supplementary Table 1**). * experimentally confirmed by TEM and ThT, ** annotation according to AmyPro (http://amypro.net/) and CPAD2.0 (https://web.iitm.ac.in/bioinfo2/cpad2). **b) Analysis of peptides derived from predicted APRs in LinB.** Seven hexapeptides derived from the APRs identified by LinB (top) were synthesised together with ten variants carrying mutations designed to suppress their aggregation (marked by red and green letters, respectively). All seven hexapeptides derived from LinBwt were experimentally verified to form fibrils (representative micrographs shown). The presence or absence of fibrils in the samples of peptides carrying designed mutations is indicated by tick mark or cross, respectively. The correspondence between the AggreProt score (aggregation threshold > 0.25) and experimental results is encoded by green colour, failed prediction of the outcome by red. **c) Results of experimental validation of AggreProt prediction using a set of 34 hexapeptides derived from AmyPro database and model protein LinB.**

Next, we experimentally validated the aggregation propensity of 17 hexapeptides selected from a subset of proteins from AmyPro database^50^. AmyPro is one of the several databases that compiles sequences of aggregation-prone proteins together with manually curated amyloidogenic sequence regions (APRs) found in literature. Noticeably, a lot of the annotated APRs are unusually long, as discussed in the *Model validation, comparison, and benchmarking* section above (longer than 50 amino acids). We therefore revisited the primary literature of several proteins from the database to investigate how the annotation has been made. One of the examples is γ-crystallin D (P07320) which is a protein found in human eye lenses and its aggregation is a hallmark of cataract^77,78^. Based on the AmyPro database, residues 1-79 are non-amyloidogenic, whereas residues 80-163 are annotated as APR (**Figure 4a**, green and red background, respectively). The annotation has been made based on the study which used pepsin digestion of the aggregates followed by MS/MS identification of the amyloid core spanning the residues 81-163 as annotated in the AmyPro^79^. The fragment was further digested in-gel using trypsin and subsequent MALDI-MS analysis enabled identification of 6 peptide fragments (80-88, 100-115, 118-140, 143-152, 154-163), suggesting that the cleavage sites are accessible by the protease. Interestingly, these fragments nicely overlap with sequence stretches with high AggreProt score (**Figure 4a**) which might indicate that they play an important role in stabilisation of the amyloid core via steric-zipper interactions. To test this, we selected two hexapeptides corresponding to the residues found in the trypsin fragments (peptide 13: 129-135, peptide 14: 121-126) with the highest AggreProt score, and two hexapeptides (peptide 16: 94-99, peptide 17: 150-155) which were not identified by the trypsin digestion and their AggreProt score is close to zero (**Figure 4a**). Additionally, we included one more hexapeptide with a high AggreProt score (peptide 15: 41-46), which is annotated as non-amyloid in the AmyPro despite experimentally characterised to form amyloid fibrils^80^. The TEM analysis of the hexapeptides after 4 weeks of incubation reveals that the AggreProt predictions were correct (**Figure 4a**) and demonstrating that the annotations in both AmyPro and CPAD2.0^49^ databases are indeed not accurate. We identified further missannotations in six additional proteins from the AmyPro database (**Supplementary Figure 10**) which included 8 false positives and 6 false negatives (**Supplementary Table 1**). This is an exceedingly important finding (though not the first one of this kind^27^) since there are numerous new aggregation prediction algorithms that use these databases for benchmarking without any further scrutiny or experimental validation^52,81–83^. Full details of each TEM micrograph obtained and their interpretation regarding the aggregating nature of the 37 analysed hexapeptides are provided in **Supplementary File 1**.

Altogether, we demonstrate successful utilisation of AggreProt in two experimental case studies. Using our newly developed algorithm, we identified seven APRs in a model enzyme and correctly assessed the biological relevance of four of them based on their solvent accessibility within the structure. Moreover, we were able to design mutations into these APRs that increase solubility and decrease the aggregation propensity of the protein. We firmly believe that we can improve these properties even further in future studies by systematically optimising the design strategy. Finally, AggreProt showed great accuracy (80 %) in identifying the aggregation propensity of isolated hexapeptides which were wrongly annotated in the databases of protein aggregation. The newly characterised 34 hexapeptide will be deposited to the WaltzDB2.0 database to expand its content and increase its impact for the further development of *in-silico* aggregation prediction tools in the future.

## Conclusions

In this study, we trained two novel protein aggregation predictors based on deep neural networks using two different feature representations of hexapeptides from WaltzDB. The predictors achieved great performance on the hexapeptide validation dataset (AuROCC scores of 0.85 and 0.89 for linear and static models, respectively). Using whole sequence validation datasets based on the AmyPro database, the performance was lower, yet still equivalent or better compared to the state-of-the-art algorithms. The lower scores can be attributed to the discrepancy between sequence length of the training (i.e., six amino acids) and validation (full sequences), small size of the training dataset (1406 entries), binary labels in the AmyPro database, and most importantly, to the incorrect annotations in the database discovered during this work. It has to be noted that recent artificially generated solubility sets may compensate for this lack of correctly annotated data in the immediate future^84^.

To truly test the performance of our predictors and to demonstrate their applicability, we carried out two experimental validations. First, we selected 34 hexapeptides identified by AggreProt in model haloalkane dehalogenase LinB and seven other proteins from AmyPro, and experimentally determined their aggregation. The experimental results matched the predictions in 79% of cases and, most importantly, confirmed our initial scepticism regarding misannotations of long (longer than 50 amino acids) APRs in the validation dataset. Based on this experience, we advise caution when relying on purely *in silico* validation of computational aggregation predictors. The newly characterised 34 hexapeptide will be deposited to the WaltzDB2.0 database to extend its content and increase its impact for further development of *in-silico* aggregation prediction tools in the future.

Finally, we showcase an example of the AggreProt use for protein design. Specifically, we correctly identified seven APRs in a model haloalkane dehalogenase LinB and correctly assessed the biological relevance of four of them based on their solvent accessibility within the structure. Moreover, we were able to design mutations into these APRs that increase solubility and decrease aggregation propensity of the protein, resulting in up to doubled soluble protein expression compared to the wild type without compromising its functionality. We firmly believe that we can improve these properties even further in future studies by systematically optimising the design strategy, making AggreProt a valuable tool for solubilizing proteins.

## Acknowledgements

The Authors are thankful to Dr. Jiri Novacek, Dr. Zuzana Hlavenkova, and Mgr. Michal Babiak at the Cryo-Electron Microscopy and Tomography Core Facility (CEITEC) for providing access and training to TEM. Computational resources were provided by the e-INFRA CZ and ELIXIR-CZ projects (90254 and LM2023055), supported by the Ministry of Education, Youth and Sports of the Czech Republic. The authors thank the RECETOX Research Infrastructure (No. LM2023069). This research was also supported by the European Union’s Horizon 2020 Research and Innovation Programme under grant agreement No. 857560 and Horizon Europe Framework Programme under the grant agreement No. 101136607 (CLARA), and by the project National Institute for Neurology Research (nr. LX22NPO5107 MEYS): Financed by European Union – Next Generation EU.

## Notes

### Competing Interest Statement

The authors have declared no competing interest.

### Summary of Updates

The text was reworked for clarity. In silico validation completed. Extended experiments and added information about variants haloalkane dehalogenase activity. Figures improved.

