## Supplementary Materials for "Deep Learning-Based Prediction and Suppression of Protein Aggregation- Prone Regions"

### Prediction of Aggregation Prone Regions in Proteins Using Deep Neural Networks and Their Suppression by Computational Design

**Vojtech Cima[1,#], Antonin Kunka[2,3,#,§], Joan Planas-Iglesias[2,3], Ekaterina Grakova[1], Martin Havlasek[2,3], Madhumalar Subramanian[2], Michal Beloch[1], Martin Marek[2,3], Katerina Slaninova[1], Jiri Damborsky[2,3], Zbynek Prokop[2,3,*], David Bednar[2,3,*], Jan Martinovic[1,*]**

[1] IT4Innovations, VSB – Technical University of Ostrava, 17. Listopadu 2172/15, 70800, Ostrava-Poruba, Czech Republic
[2] Loschmidt Laboratories, Department of Experimental Biology and RECETOX, Faculty of Science, Masaryk University, Kotlarska 2, Brno, Czech Republic
[3] International Clinical Research Centre, St. Anne's University Hospital, Pekarska 53, Brno, Czech Republic
§The current affiliation of the Author is: Protein Biophysics Group, Department of Biotechnology and Biomedicine, Technical University of Denmark, Søltofts Plads, Building 227, 2800, Kgs. Lyngby, Denmark

#### Appendix A. Supplementary data

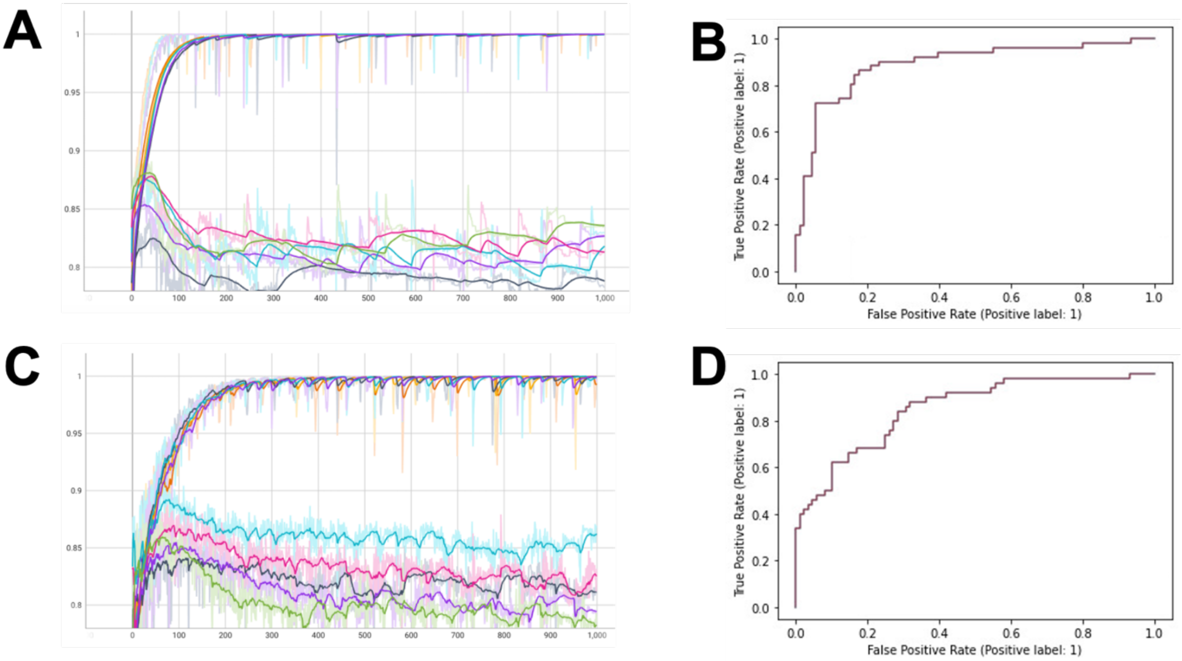

**Supplementary Figure 1: Training, validation and testing of the linear and static models on hexapeptide data.** Panels **A** and **C** show the results of the hyperparameter optimisation procedure (training and validation) of the linear and static models, respectively on WaltzDB-90 dataset. Each data series (different colours) represents each of the data-splits (1:4) produced during the 5-fold cross-validation. On the x-axis is represented the training epoch, and on the y-axis the corresponding AuROCC value. The upper lines correspond to the training data (max(AuROCC) = 1) and the lower ones to the validation data, where the max(AuROCC) is of 0.88 and 0.89 for the linear (**A**) and static models (**C**), respectively. Panels **B** and **D** show the results, in the form of a ROC curve depicted as a red line, of the testing on the independent set WaltzDB-10 for the linear (AuROCC = 0.89) and static (AuROCC = 0.85) models, respectively.

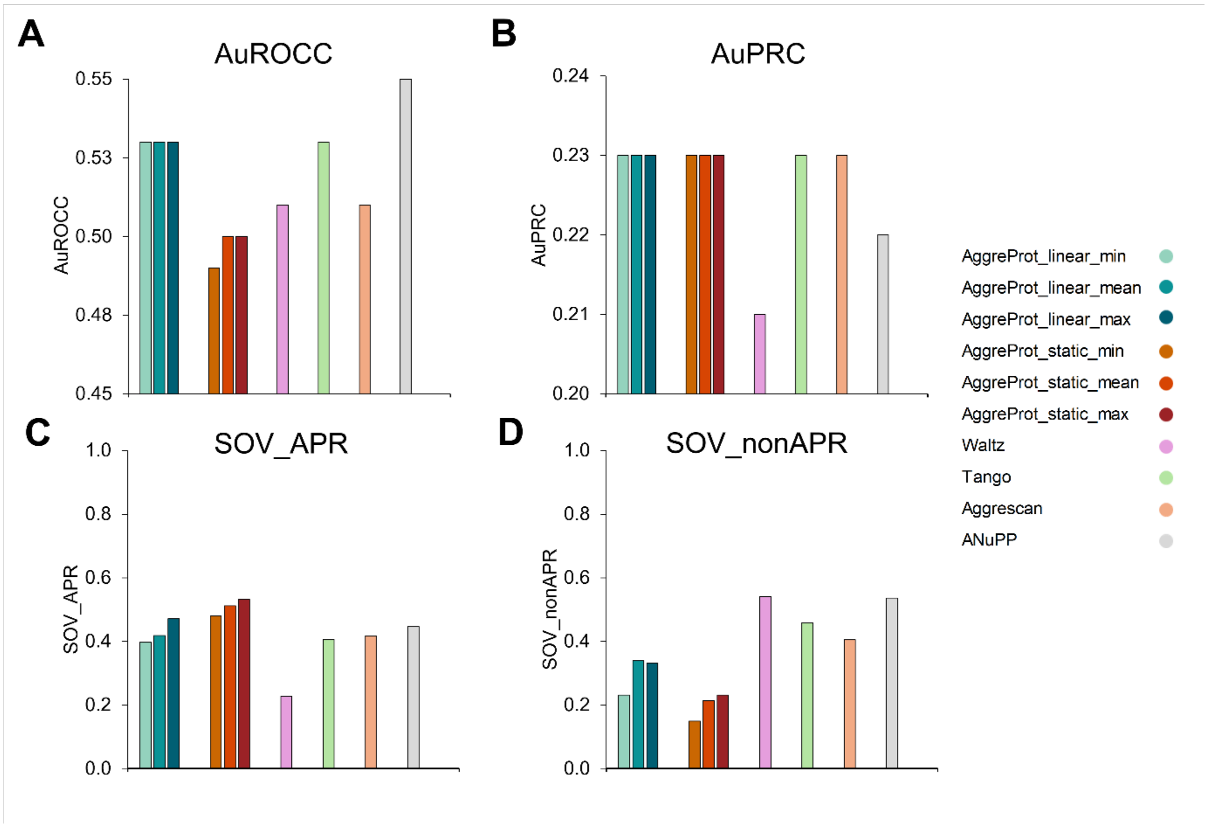

**Supplementary Figure 2: Performance of the trained networks using different aggregating metrics and other state-of the art predictors on AmyPro37.** Different evaluation metrics are shown for the performance of the linear (blue hues) and the static (red hues) predictors on AmyPro37, along with Waltz (pink), Tango (green), Aggrescan (orange), and ANuP (grey). **A**: Area under the Receiver Operating Characteristic Curve (AuROCC). **B**: Area under the Precision-Recall Curve (AuPRC). **C**: Averaged Segment OVerlap score for Aggregation Prone Regions (SOV_APR). **D**: Averaged Segment OVerlap score for Non-Aggregation Prone Regions (SOV_nonAPR). The magnitude of each metric is shown in the y axis.

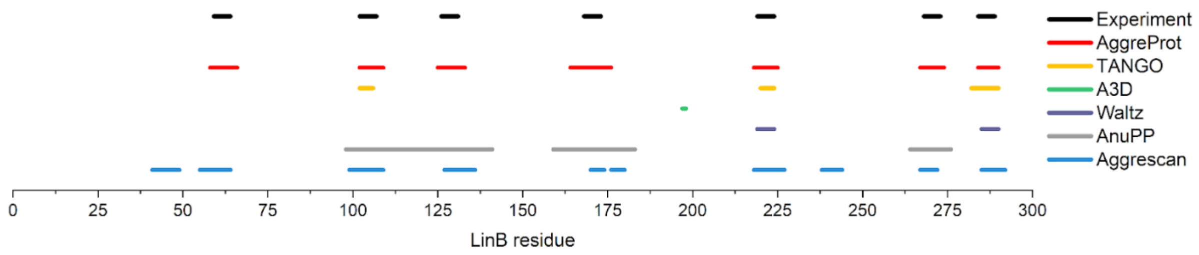

**Supplementary Figure 3: Comparison of APR prediction in LinB based on different algorithms.** The experimental results correspond to the hexapeptides whose aggregation was determined in this study. The APR annotations were made using default settings of each predictor according to authors recommendations.

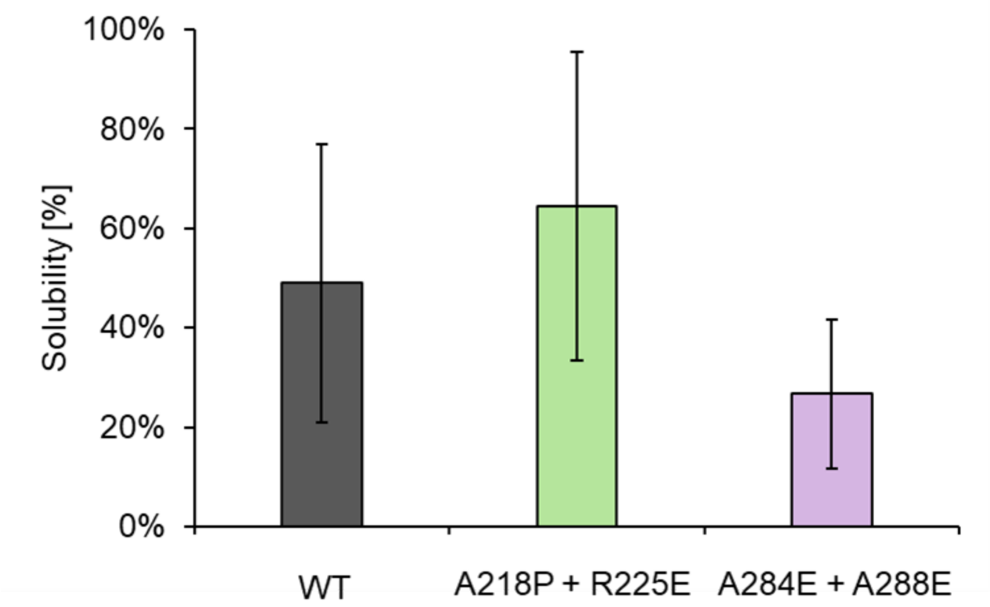

**Supplementary Figure 4: Solubility determination using SDS-PAGE for LinBwt and two best variants.** Poor reproducibility of this experiment resulted in high deviations, therefore soluble yield was selected as better metric for solubility comparisons.

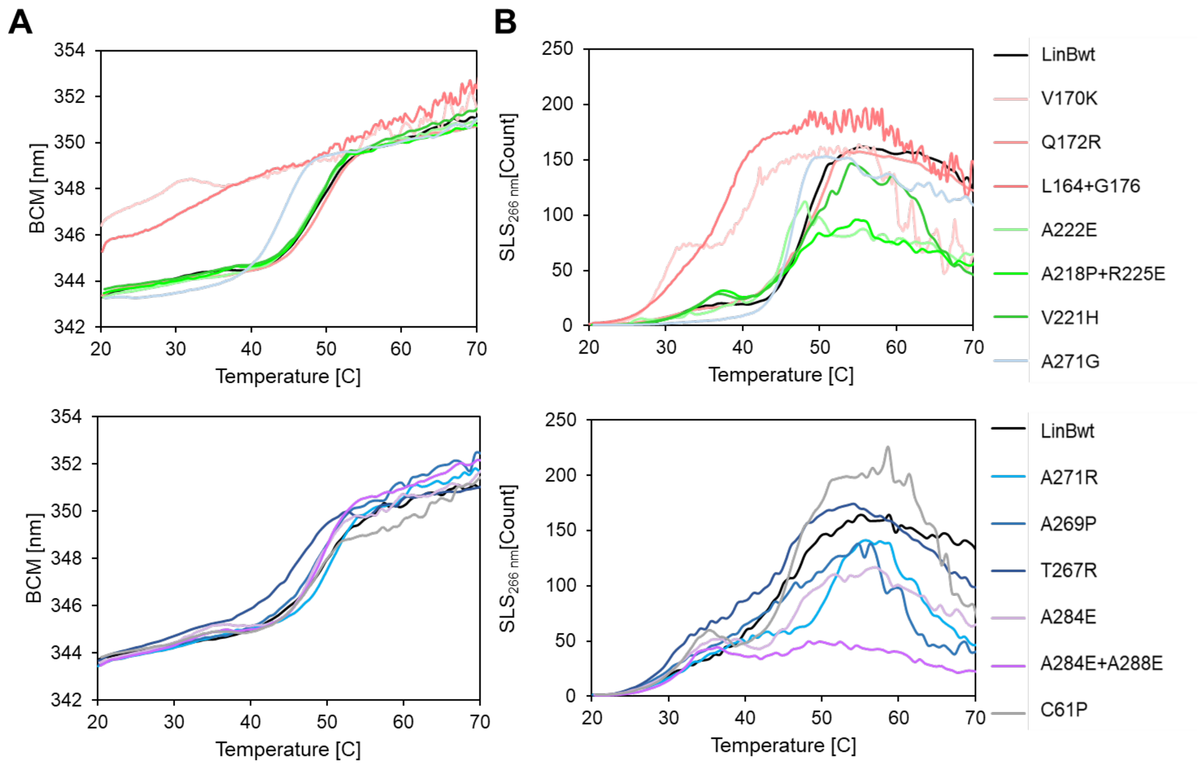

**Supplementary Figure 5: Temperature scanning experiments showing unfolding using fluorescence measurements (A) and aggregation measured using SLS (B).** Monitoring of unfolding (barycentric mean of fluorescence -BCM) and aggregation ($SLS_{266nm}$) aer shown as a function of temperature for LinBwt and different mutants. The traces shown in different colour hues represent different protein variants, indicated in the right-side legend.

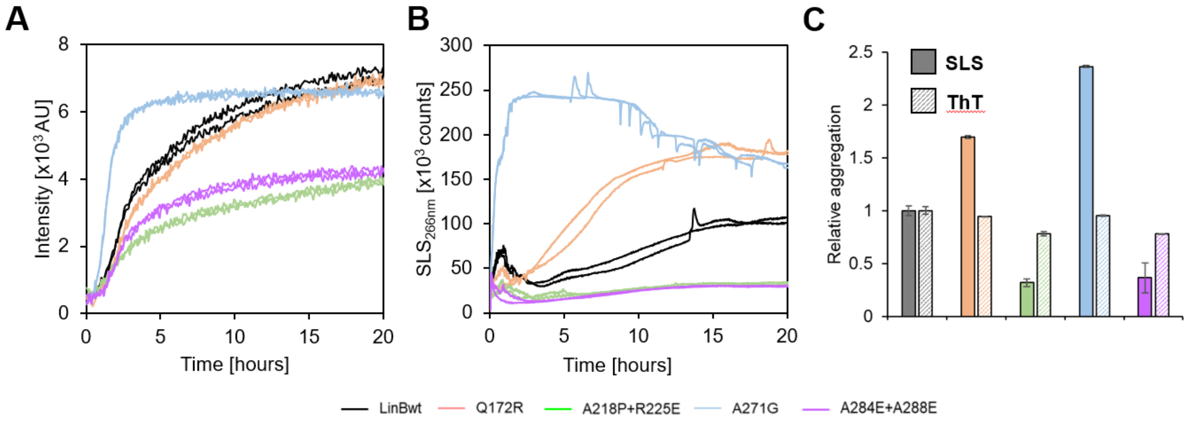

**Supplementary Figure 6: Aggregation kinetics measured using ThT (A) and SLS (B).** The aggregation of respective LinB variants was monitored at 37°C for 24h. Despite similar shapes and initial slopes of the curves, the amplitudes that correspond to the final amounts of aggregates differ **(C)**, confirming the results of previous scanning indicating reduced aggregation for A218P+R225E and A284E+A288E mutants.

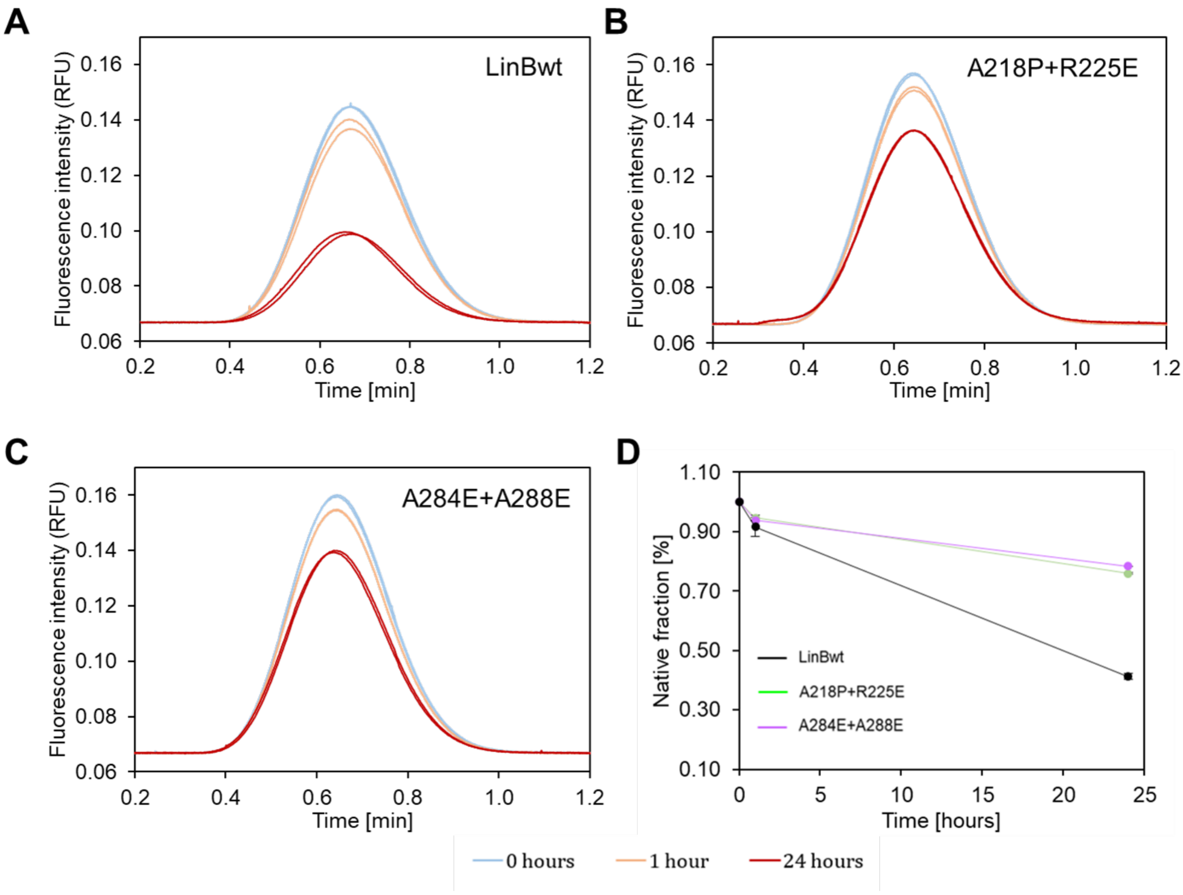

**Supplementary Figure 7: Aggregation of LinBwt and selected variants measured by flow-induced dispersion analysis (FIDA). Taylorgrams of A)** (LinB WT), **B)** LinB A218P+R225E, and **C)** LinB A284E+A288E corresponding to the soluble protein fraction upon 0 (blue), 1 (orange) and 24 (red) hours of incubation at 37°C. All measurements were done in duplicate. **D) Relative concentration of soluble fractions** obtained by the integration of the Taylorgrams in A, B, and C, as a function of time.

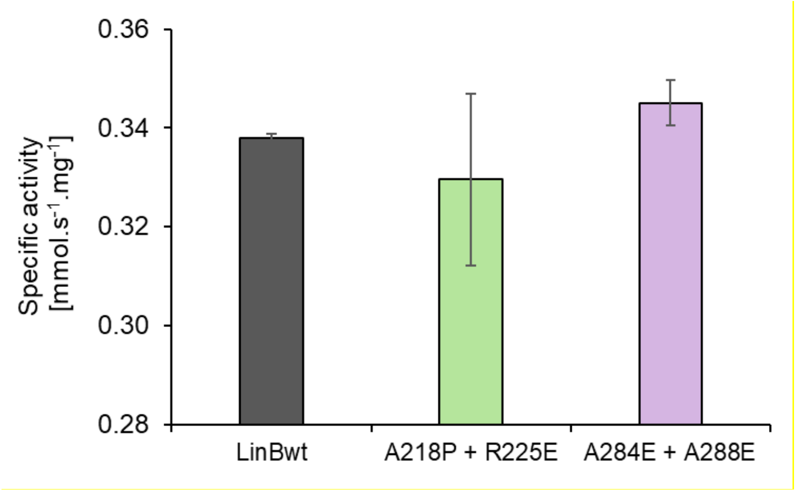

**Supplementary Figure 8: Specific activities towards 1,2-dibromoethane.** The experiments were conducted at 37 °C (see Method section for further details).

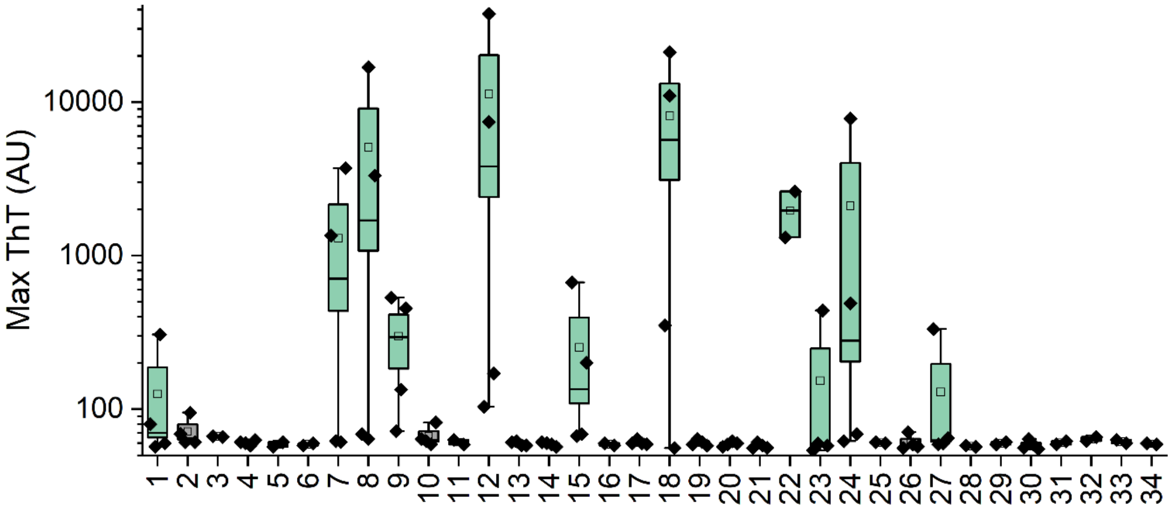

**Supplementary Figure 9. Aggregation of hexapeptides monitored by ThT.** Maximum intensity of ThT fluorescence (black symbols) of peptides (numbering shown on the x-axis corresponds to the naming in supplementary table 1) incubated at 1 mM for one month (see methods for details). The experiments were done in duplicates or quadruplicates in case of samples that had to be dissolved using DMSO or HFIP as stated in the Supplementary Table 1. The samples were considered ThT positive when the mean fluorescence intensity (squares) was above 100 AU. The box, vertical line and whiskers indicate 1 SE, median value, and outliers, respectively.

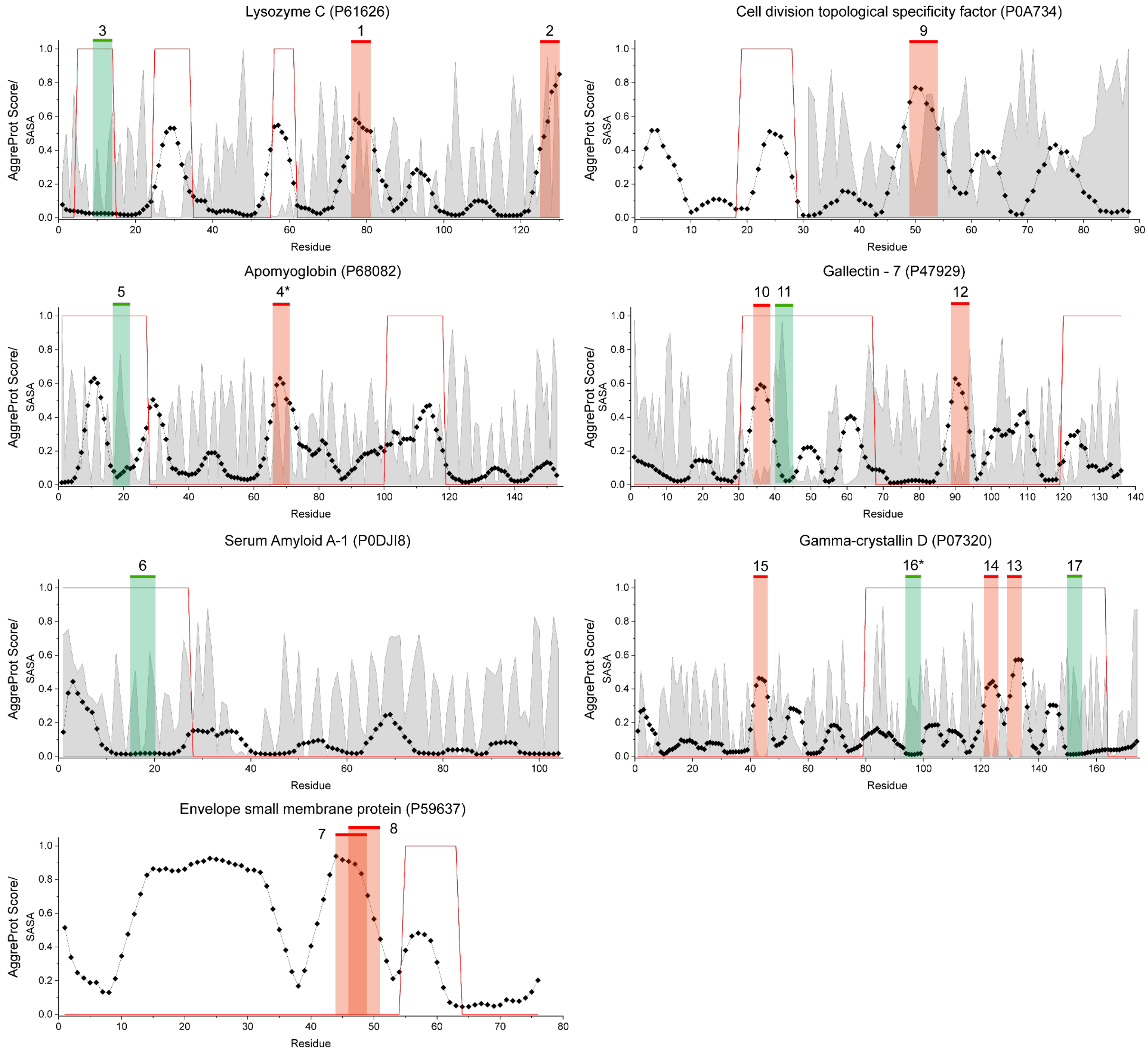

**Supplementary Figure 10: Experimental validation of AggreProt on hexapeptides derived from AmyPro37 dataset.** Aggregation propensity of seven proteins (residues represented in the x-axis) according to the AggreProt prediction (y-axis) is depicted (dotted, solid black line profile). SASA (using the same propensity y-axis) is shown in grey shade. The red line shows AmyPro annotation: it takes a value of 0 and 1 if the region is predicted as non-APR or APR, respectively. The different hexapeptides selected was experimental evaluation are indicated by shaded coloured areas with a top solid thick line (red and green). Green hexapeptides correspond to cases that were originally classified as false negative predictions but are actually correct negative predictions, i.e., identified as non-APR by AggreProt, and experimentally confirmed as non-APRs (except #16 for which results were inconclusive), although annotated as APR in the AmyPro database. The red hexapeptides correspond to either (i) false positives (i.e., annotated as non-APRs in the AmyPro database, classified as APRs by AggreProt, and experimentally verified to be APRs, #1, 2, 4, 7, 8, 9, 12, and 15), or (ii) true positives (annotated as APRs in the AmyPro database, classified as APRs by AggreProt, and experimentally verified to be APRs, #10, 13, and 14).

**Supplementary Table 1: Overview of the experimentally characterized hexapeptides.** Peptides 1-17 represent a selection of hexapeptides derived from AmyPro37 dataset that were predicted as false positive (FP), false negative (FN), or true positives (TP). Peptides 18-34 were derived from LinB APRs and their mutants. AP score refers to the AggreProt score of the respective peptide and Solub. agent corresponds to the solvent used to dissolve the peptide. The columns TEM and ThT qualitatively describe whether the peptide aggregates significantly (++), slightly (+), or no aggregates were detected (-) by the two methods. Exp. Agg. conclude the experimentally observed aggregation (Y-yes, N-no).Pred vs. Exp. describes the consensus between the predicted and experimental results (Y for Yes, N for No, and N.D. for the cases where we could not unequivocally experimentally determine the aggregation behaviour of the hexapeptide). The raw experimental data supporting our classification are shown in **Supplementary File 1**.

| #P | Protein | Sequence | Type | AP score | Solub. agent | TEM | ThT | Exp. Agg. | Pred. Vs Exp. |
|---|---|---|---|---|---|---|---|---|---|
| 1 | Lysozyme C | ACHLSC | FP | 0.530 | HFIP | + | + | Y | Y |
| 2 | | VQGCGV | FP | 0.640 | HFIP | + | - | Y | Y |
| 3 | | ARTLKR | FN | 0.025 | buffer | - | - | N | Y |
| 4 | Apomyoglobin | TVVLTA | FP | 0.548 | DMSO | - | - | amorphous | N.D. |
| 5 | | VEADIA | FN | 0.077 | buffer | - | - | N | Y |
| 6 | Serum Amyloid A | RDMWRA | FN | 0.017 | buffer | - | - | N | Y |
| 7 | Envelope small membrane protein | CNIVNV | FP | 0.866 | HFIP | + | + | Y | Y |
| 8 | | IVNVSL | FP | 0.725 | DMSO | ++ | + | Y | Y |
| 9 | Cell division topological specificity factor | VICKYV | FP | 0.678 | HFIP | + | + | Y | Y |
| 10 | Galectin-7 | HVNLLC | TP | 0.513 | HFIP | + | - | Y | Y |
| 11 | | GEEQGS | FN | 0.087 | buffer | - | - | N | Y |
| 12 | | VLIIAS | FP | 0.505 | DMSO | + | + | Y | Y |
| 13 | Gamma-crystallin D | SWVLYE | TP | 0.497 | DMSO | ++ | - | Y | Y |
| 14 | | IHSLNV | TP | 0.393 | DMSO | ++ | - | Y | Y |
| 15 | | GCWMLY | FP | 0.407 | HFIP | + | + | Y | Y |
| 16 | | EREDYR | FN | 0.014 | buffer | n.d. | - | inconclusive | N.D. |
| 17 | | DYRRYQ | FN | 0.013 | DMSO | - | - | N | Y |
| 18 | LinBwt | IACDLI | n.a. | 0.520 | HFIP | ++ | + | Y | Y |
| 19 | | VLVVHD | n.a. | 0.429 | DMSO | + | - | Y | Y |
| 20 | | QGIAYM | n.a. | 0.358 | DMSO | + | - | Y | Y |
| 21 | | VFVEQV | n.a. | 0.511 | DMSO | + | - | Y | Y |
| 22 | | DVVAIA | n.a. | 0.368 | buffer | ++ | + | Y | Y |
| 23 | | VAGAHF | n.a. | 0.359 | DMSO | + | - | Y | Y |
| 24 | | AAIAAF | n.a. | 0.425 | DMSO | + | + | Y | Y |
| 25 | LinB174 (APR1) | IAPDLI | n.a. | 0.084 | buffer | + | - | Y | N |
| 26 | LinB161 (APR4) | VFKEQV | n.a. | 0.117 | DMSO | + | - | Y | N |
| 27 | LinB162 (APR4) | VFVERV | n.a. | 0.304 | DMSO | + | + | Y | Y |
| 28 | LinB164 (APR5) | DVVEIA | n.a. | 0.305 | buffer | + | - | Y | Y |
| 29 | LinB166 (APR5) | DVHAIA | n.a. | 0.281 | buffer | ++ | - | Y | Y |
| 30 | LinB167 (APR6) | VAGGHF | n.a. | 0.332 | DMSO | - | - | N | N |
| 31 | LinB168 (APR6) | VAGRHF | n.a. | 0.246 | buffer | - | - | N | Y |
| 32 | LinB171 (APR7) | AADAAF | n.a. | 0.040 | buffer | + | - | Y | N |
| 33 | LinB172 (APR7) | EAIAAF | n.a. | 0.275 | buffer | ++ | - | Y | Y |
| 34 | LinB173 (APR7) | EAIAEF | n.a. | 0.143 | buffer | + | - | Y | N |

**Supplementary File 1: Overview of TEM micrographs and their analysis**. First (Protein) and second (Peptide Sequence) columns describe the protein to which the hexapeptide belongs and its sequence, respectively. The third column, AggreProt,  describes the reason for the peptide selection: FP (False Positive) denotes a peptide predicted as aggregating by AggreProt but not recorded as such in AmyPro. FN (False Negative) denotes a peptide predicted as non-aggregating by AggreProt but corresponding to an aggregation described in AmyProt. TP (True Positive) and TN (True Negative) denote an AggreProt prediction matching the data recorded in AmyProt, either aggregating (TP) or non-aggregation (TN). AggreProt AV (Average Value) denotes the average of AggreProt predictions for each residue in the hexapeptide. Solubilization denotes the solubilizing agent: either 'buffer' (50 mM sodium phosphate, pH 7.4), 'DMSO' (Dimethylsulfoxid) or 'HFIP' (hexafluoroisopropanol). Most aggregated column displays micrographs showing the regions of the sample with clearer fibrils formation, and Least aggregated column displays micrographs of the most solubilized regions observed. The column other structures shows evidence of formation of other amorphous or aggregation structures or hydrogels. Aggregation +/- denotes our interpretation of the micrographs: '+' indicating aggregation presence and '-' indicating aggregation absence. Finally, we include qualitative observation notes in the last column Notes.
